# Introgression across evolutionary scales suggests reticulation contributes to Amazonian tree diversity

**DOI:** 10.1101/2019.12.12.873927

**Authors:** Rowan J. Schley, R. Toby Pennington, Oscar Alejandro Pérez-Escobar, Andrew J. Helmstetter, Manuel de la Estrella, Isabel Larridon, Izai Alberto Bruno Sabino Kikuchi, Timothy Barraclough, Félix Forest, Bente Klitgård

## Abstract

Hybridization has the potential to generate or homogenize biodiversity and is a particularly common phenomenon in plants, with an estimated 25% of species undergoing inter-specific gene flow. However, hybridization in Amazonia’s megadiverse tree flora was assumed to be extremely rare despite extensive sympatry between closely related species, and its role in diversification remains enigmatic because it has not yet been examined empirically. Using members of a dominant Amazonian tree family (*Brownea*, Fabaceae) as a model to address this knowledge gap, our study recovered extensive evidence of hybridization among multiple lineages across phylogenetic scales. More specifically, our results uncovered several historical introgression events between *Brownea* lineages and indicated that gene tree incongruence in *Brownea* is best explained by introgression, rather than solely by incomplete lineage sorting. Furthermore, investigation of recent hybridization using ∼19,000 ddRAD loci recovered a high degree of shared variation between two *Brownea* species which co-occur in the Ecuadorian Amazon. Our analyses also showed that these sympatric lineages exhibit homogeneous rates of introgression among loci relative to the genome-wide average, implying a lack of selection against hybrid genotypes and a persistence of hybridization over time. Our results demonstrate that gene flow between multiple Amazonian tree species has occurred across temporal scales, and contrasts with the prevailing view of hybridization’s rarity in Amazonia. Overall, our results provide novel evidence that reticulate evolution influenced diversification in part of the Amazonian tree flora, which is the most diverse on Earth.

## Introduction

Reproductive isolation is often seen as a prerequisite for speciation and as a defining feature of species (Barraclough 2019; Mayr 1942). Despite this, hybridization between species is known to occur and has several different outcomes, from the erosion of evolutionary divergence (Kearns *et al*., 2018; Vonlanthen *et al*., 2012) to the formation of entirely new ‘hybrid species’ (Mallet 2007). Neotropical rainforests harbour the highest levels of plant diversity on Earth (Antonelli & Sanmartín 2011), and to date there has been little convincing evidence that hybridization consistently occurs between tree species therein. Indeed, the prevailing view has been that hybridization between tropical tree species is an exceptionally rare event (Ashton 1969; Ehrendorfer 1970; Gentry 1982). Although a few observations of reproductive biology support this, such as the intersterility between lineages of *Inga*, a species-rich genus in the legume family (Koptur 1984), evidence for hybridization has been poorly tested empirically in tropical floras. The only potential example of hybridization among neotropical trees which has been documented using DNA sequence data involves two species of *Carapa* (Scotti-Saintagne *et al*., 2013) found in Amazonia, the largest expanse of rainforest in the world, containing at least 6,800 tree species (Cardoso *et al*., 2017).

One factor that suggests hybridization might occur more frequently in neotropical rainforest tree species, if their reproductive isolation is not absolute, is the remarkable level of sympatry found for closely related species. In several neotropical lineages, including *Inga, Guatteria* and *Swartzia*, many recently diverged species co-occur (Dexter *et al*., 2017), often to a remarkable degree. One such example of this is the co-occurrence of 19 *Inga* species in a single hectare of the Ecuadorian Amazon (Valencia, Balslev, & Miño 1994). As such, for many rainforest taxa the opportunity for hybridization is constantly present.

Hybridization can have a range of evolutionary consequences. In many cases, hybridization simply results in the formation of sterile, maladapted offspring with poor reproductive fitness, allowing genetic isolation between species to be maintained (i.e., reinforcement (Hopkins 2013)). In other cases, there may be a permanent movement of genetic material from one lineage to another (Rieseberg & Wendel 1993), which is known as ‘introgression’. This transfer of genetic material through hybridization may confer a selective advantage to resultant offspring (Taylor & Larson 2019), in which case it is referred to as ‘adaptive introgression’. Adaptive introgression is most often observed between closely-related taxa during the invasion of new habitats (Suarez-Gonzalez, Lexer, & Cronk 2018; e.g. Whitney, Randell, & Rieseberg 2010). Furthermore, hybridization can lead to rapid evolutionary radiations. This occurs through the re-assembly of standing genetic variation which has accumulated between diverging lineages, and which has already been subject to selection. This ‘combinatorial’ process is much more rapid than the gradual accumulation of variation through mutation, and the passage of these variants through hybridization often fuels rapid diversification events (Marques, Meier, & Seehausen 2019). This has been demonstrated in a wide range of taxa, including sunflowers, African cichlid fish and Darwin’s finches (Lamichhaney *et al*., 2018; Meier *et al*., 2017; Rieseberg *et al*., 2003).

Introgression can occur at different rates in different regions of the genome (Payseur 2010). Regions under divergent selection may remain distinct due to reduced fitness of hybrid genotypes at such loci, resulting in a low rate of introgression. Conversely, regions under little or no selection tend to introgress more freely, becoming homogenised between species. Moreover, if there is selection for hybrid genotypes (as in adaptive introgression), the rate of introgression for a region may be further increased relative to the rest of the genome (Gompert, Parchman, & Buerkle 2012). Such a process has been demonstrated in temperate-zone tree species, where divergence between hybridizing lineages is maintained through environmentally-driven selection (Hamilton & Aitken 2013; Sullivan *et al*., 2016). This explains why species that hybridize can remain as biologically distinct (and taxonomically identifiable) groups despite undergoing genetic exchange with other species (Abbott *et al*., 2013; Seehausen *et al*., 2014).

*Brownea*, a member of the legume family (Fabaceae), is a characteristic tree genus of lowland neotropical rainforests and contains around 27 species distributed across northern South America. Previous work indicates that there is a broad degree of phylogenetic incongruence evident in this genus (Schley *et al*., 2018) which might indicate hybridization, although this could also result from incomplete lineage sorting. However, there are numerous *Brownea* hybrids in cultivation (e.g. *B. x crawfordii* (Crawford & Nelson 1979) and *B. hybrida* (Backer 1911)), as well as several instances of putative hybridization among *Brownea* lineages in the wild. The most notable of these instances is that proposed between the range-restricted *Brownea jaramilloi* (Pérez *et al*., 2013) and the wide-ranging *B. grandiceps*, which co-occur in the Ecuadorian Amazon (Fig. 1). There are multiple morphological distinctions between these two species, including differences in inflorescence colour and structure, growth habit, tree height and leaf morphology (Pérez *et al*., 2013). Although they co-occur, these species favour different habitats: *B. jaramilloi* grows on ridge tops and hillsides, whereas *B. grandiceps* shows a slight preference for swamps and valleys but is more evenly distributed (pers. obs. & Klitgaard 1991; Pérez *et al*., 2013). Despite this, hybridization appears to occur, as evidenced by the existence of a putative hybrid between these two species known as *B*. “rosada” (Fig. 1). *Brownea “*rosada*”* displays an intermediate morphology between its two parental species, producing pink flowers. The hypothesis of a *B. jaramilloi* x *B. grandiceps* hybrid has not yet been tested, however, using molecular data.

**Figure 1:**
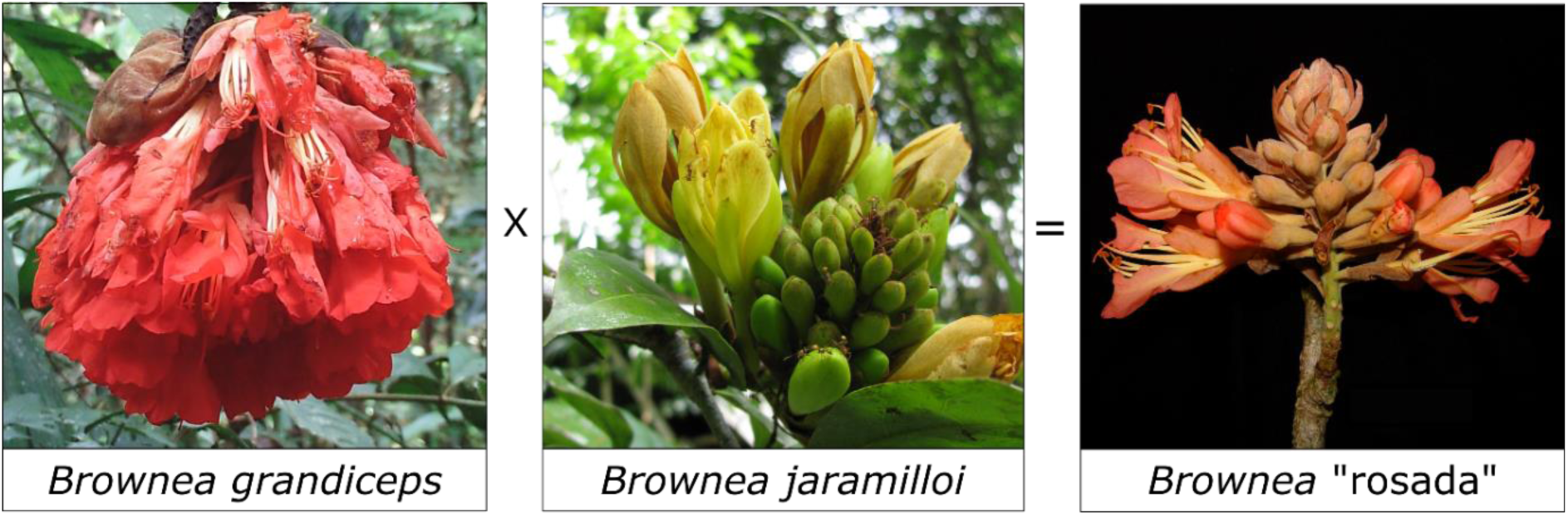
Two co-occurring *Brownea* lineages (*Brownea grandiceps* (photograph © Rowan Schley) and *Brownea jaramilloi* (photograph © Xavier Cornejo)) as well as their putative hybrid *Brownea* “rosada” (photograph © J. L. Clark).

As a member of the legume family, which dominates Amazonian forests (ter Steege *et al*., 2013), and with its apparent propensity for hybridization, *Brownea* is an excellent system with which to study the phylogenetic patterns and genomic architecture of introgression in neotropical rainforest trees.

Systematically documenting hybridization at a range of time scales and taxonomic levels within this group could reveal how admixture has contributed to the assembly of one of the world’s richest floras. Accordingly, this investigation aims to answer the following questions:

1. Is there evidence of introgression at a deep phylogenetic level in these lineages of neotropical rainforest tree?
2. Is there evidence of more recent gene flow, and if so, does this occur evenly across most of the genome?

## Materials and Methods

### Phylogenetic analysis

Sequences from 226 nuclear genes were used to elucidate the evolutionary relationships between *Brownea* species using a phylogenomic approach. To do so, 23 of 27 lineages were sampled within *Brownea*, including the three subspecies of *B. coccinea* and *B*. “rosada”, a putative hybrid lineage of *B. grandiceps* and *B. jaramilloi*. In total, 59 accessions were sampled within *Brownea*, as well as an additional 13 outgroup taxa from the genera *Macrolobium, Heterostemon, Paloue* and *Browneopsis*, which form part of the ‘Brownea clade’ (Fabaceae, subfamily Detarioideae (LPWG, 2017)). The list of accessions and their associated information can be found in Supporting Information Table S1.

DNA sequence data were generated using leaf material collected from herbarium specimens and silica-gel dried accessions. Genomic DNA was extracted using the CTAB method (Doyle & Doyle 1987), and sequencing libraries were prepared using the NEBNext® Ultra™ II DNA Kit (New England Biolabs, Massachusetts, USA) with a modified protocol to account for fragmented DNA, and including a ∼600bp size-selection step. Hybrid bait capture was performed using the MyBaits protocol (Arbor Biosciences, Michigan, USA) to target 289 phylogenetically-informative nuclear genes, using a bait kit designed for the legume subfamily Detarioideae within which *Brownea* is nested (Ojeda *et al*., 2019). The final library pools were sequenced either on the Illumina MiSeq platform (Illumina, San-Diego, USA) at RBG Kew, or on the Illumina HiSeq platform by Macrogen Inc. (Seoul, South Korea).

DNA sequencing reads were quality-checked with the program *FASTQC* v0.11.3 (Andrews 2010), and were subsequently trimmed using *Trimmomatic* v.0.3.6 (Bolger, Lohse, & Usadel 2014) in order to remove adapter sequences and to quality-filter reads. *Trimmomatic* settings permitted <4 mismatches, a palindrome clip threshold of 30 and a simple clip threshold of 6. Bases with a quality score <28 and reads shorter than 36 bases long were removed from the dataset. Following quality-filtering, loci were assembled using *SPAdes* v3.11.1 (Bankevich *et al*., 2012) by the HybPiper pipeline v1.2 (Johnson *et al*., 2016), and potentially paralogous loci were removed using the Python (Python Software Foundation 2010) script ‘*paralog_investigator*.*py*’, distributed with the HybPiper pipeline. All sequences were aligned by gene region using *MAFFT* v7.215 (Katoh & Standley 2013). In order to infer gene trees for phylogenetic network analysis, the 226 recovered gene regions were further refined to include only 20 taxa, representing a single accession per lineage in *Brownea*, using *Macrolobium colombianum* as the outgroup taxon. Where applicable, samples were chosen by comparing the sequence recovery of conspecific accessions and choosing the individual with the best gene recovery. This resulted in 220 single-accession-per-lineage gene alignments. Further details of phylogenomic sampling, labwork, sequencing and data processing are detailed in Supporting Information Methods S1.

Gene trees were generated for each of the 226 gene alignments using *RAxML* v.8.0.26 (Stamatakis 2014) with 1,000 rapid bootstrap replicates and the GTRCAT model of nucleotide substitution, which is a parameter-rich model and hence is most suitable for large datasets. *Macrolobium colombianum* was used to root both the full and single-accession dataset analyses since it was the *Macrolobium* accession that was present in the most gene alignments. The gene trees from both datasets were used to generate species trees under the multi-species coalescent model in the heuristic version of *ASTRAL* v.5.6.1 (Zhang *et al*., 2018) using the default parameters. Monophyly was not enforced for individuals belonging to the same species (the ‘*-a*’ option in *ASTRAL*). Finally, discordance between gene trees was calculated and visualised for the full dataset using the Java program *PhyParts* v0.0.1 (2015) (https://bitbucket.org/blackrim/phyparts). The pattern of incongruence between gene trees for each node was then mapped onto the *ASTRAL* species tree using the Python script *PhyPartsPieCharts* v1.0 (https://github.com/mossmatters/MJPythonNotebooks).

### Inferring ancient introgression

Phylogenetic networks were inferred for 220 gene trees from the single-accession-per-lineage dataset to understand whether introgression occurred over *Brownea*’s evolutionary history. Networks were inferred with the program *SNaQ!*, implemented in the *Julia* v0.6.4 (Bezanson *et al*., 2017) package *PhyloNetworks* v0.11.0 (Solís-Lemus, Bastide, & Ané 2017). This program facilitates testing of whether the observed incongruence between gene trees is better explained by a model describing only ILS or by a model describing introgression. Networks were estimated by calculating quartet concordance factors (CF) for each node from the single-accession-per-lineage gene trees, since *SNaQ!* requires that each tip of the phylogenetic trees represent a single lineage. The network with the number of hybridization events (*h*) best describing the data was chosen using negative log pseudolikelihood comparison. Finally, the observed gene tree CFs were compared with those of the best-fit phylogenetic network (i.e., the model including hybridization) and those expected under a coalescent model (i.e., a ‘tree-like’ model which accounts for incomplete lineage sorting but not hybridization). The fit of the observed gene tree topologies to either the ‘network-like’ or the ‘tree-like’ model was assessed using the Tree Incongruence Checking in R (TICR) test (Stenz *et al*., 2015). This was done with the function ‘*test*.*one*.*species*.*tree’* in the R v3.4.4 (R Development Core Team 2013) package *PHYLOLM* (Ho & Ané 2014). Proportions of genes contributed between lineages by hybridization events were taken from the ‘Gamma value’ output of *PhyloNetworks*. Further details of phylogenetic network analysis are contained in Supporting Information Methods S1.

### Population-level introgression

Having examined the degree of historical reticulation among *Brownea* lineages, population genomic data were generated with ddRAD sequencing and used to investigate recent introgression at a finer taxonomic scale. More specifically, the degree of shared genetic variation was visualised for individuals of the ecologically divergent species *B. grandiceps* and *B. jaramilloi*, which co-occur in the Ecuadorian Amazon and putatively hybridize in the wild. Following this, in order to make inferences about the potential evolutionary significance of recent introgression, the rates at which different loci introgress relative to the rest of the genome were estimated for these taxa using genomic clines. In order to do this, 171 specimens in total were genotyped using ddRADseq (Peterson *et al*., 2012). Sampling consisted of 128 individuals of *Brownea grandiceps*, 40 individuals of *B. jaramilloi* and three individuals of *B*. “rosada” (the putative hybrid of *B. grandiceps* and *B. jaramilloi*), representing the relative abundance of each species in the forest plot from which they were sampled. Leaf material was collected from *Brownea* trees in the Yasuní National Park 50ha forest plot in Napo, Ecuador, and dried in a herbarium press. The sample list is shown in Supporting Information Table S2.

Genomic DNA was extracted from dried leaf material with the CTAB method and purified using a QIAGEN plant minikit (QIAGEN, Hilden, Germany) column cleaning stage. Sequencing libraries were prepared by digesting the DNA template using the restriction enzymes *EcoRI* and *mspI* (New England BioLabs, Massachusetts, USA), in accordance with the ddRADseq protocol. Samples were then ligated to universal Illumina P2 adapters and barcoded using 48 unique Illumina P1 adapters. Samples were pooled and size-selected to between 375-550 bp using a Pippinprep electrophoresis machine (Sage Science, Massachusetts, USA). Following amplification and multiplexing, sequencing was performed using a single-lane, paired-end 150bp run on the HiSeq 3/4000 platform.

DNA sequencing reads from the ddRADseq genotyping were processed *de novo* (i.e., without the use of a reference genome) using the Stacks pipeline v2.1 (Catchen *et al*., 2011). Reads were quality filtered using Stacks by removing reads which were of poor quality (i.e., had a phred score <10), following which ‘stacks’ of reads were created, and SNPs were identified among all *de novo* ‘loci’ and across individuals. This was done using a minimum coverage depth (the ‘*-m’* flag in Stacks) of three and a within-individual mismatch distance (*-M*) of seven nucleotides. Individuals with sequencing coverage under 7.5x were removed. Loci found in fewer than 40% of individuals and sites with a minor allele frequency threshold of 5% were filtered out using the *‘populations*’ module of Stacks to account for genotyping error. Model parameters for the various programs within the Stacks pipeline were chosen using the recommendations in Paris *et al*. (2017), as well as through running the pipeline with multiple different parameter settings. This resulted in a dataset containing 22,046 loci with 120,085 SNPs for 171 individuals. A dataset containing only one SNP per locus was also extracted using the Stacks *populations* module for use in analyses which assumed no linkage disequilibrium. This subsetting resulted in a dataset containing 19,130 loci with 19,130 SNPs for 171 individuals. Details of population genomic sampling, library preparation, sequencing and data filtering are shown in Supporting Information Methods S2.

In order to understand the patterns of introgression at the population level, the full ddRAD dataset was used to visualise the degree of shared variation between *B. grandiceps* and *B. jaramilloi*. This was performed using principal component analysis implemented in R v3.4.4, followed by plotting with *ggplot2* (Wickham 2016), and by producing a neighbour net plot in *SPLITSTREE* v4.14.6 (Huson & Bryant 2005). Furthermore, the single-SNP-per-locus dataset containing 19,130 RAD loci was used to infer population structure with the program *fastSTRUCTURE* v1.0 (Raj, Stephens, & Pritchard 2014). The number of populations (*K*) was chosen using the value which provided the largest improvement in marginal likelihood. In addition, a *fastSTRUCTURE* run incorporating 40 random individuals from each species was performed to account for any bias which may have been incurred by differences in sample size. Finally, *NEWHYBRIDS* v1.1 (2002) (https://github.com/eriqande/newhybrids) was used to categorise individuals into different hybrid classes, using three runs on a subset of 500 randomly selected ddRAD loci due to the computational limits of the program. Runs were performed with 50,000 MCMC sweeps following 50,000 burn-in sweeps under the default parameters of the program.

The relative ‘rate’ and ‘direction’ of introgression for each locus between the two *Brownea* species was estimated using Bayesian estimation of genomic clines (*bgc*) v1.03 (Gompert & Buerkle 2011). For each locus, *bgc* compares the probability of ancestry at the locus relative to an individual’s genome wide ancestry, thereby allowing it to estimate two parameters for each locus. These parameters are α, which roughly equates to the ‘direction’ of introgression, and β, which may be summarised as the ‘rate’ of introgression for a locus (Gompert & Buerkle 2011; Gompert *et al*., 2012). In order to estimate these parameters from the single-SNP dataset consisting of 19,130 RAD loci, 50,000 MCMC generations with a 50% burn-in were used in *bgc*. Admixture proportions (i.e., mean *Q* values) generated by *fastSTRUCTURE* were used to assign each individual to three populations (*Brownea grandiceps, B. jaramilloi* and admixed). Convergence was checked for the MCMC output from *bgc* in Tracer v1.6 (Rambaut *et al*., 2015) and with the R package *coda* (Plummer *et al*., 2006) using Geweke’s diagnostic (Geweke 1991). Loci with significant ‘excess ancestry’ were identified by ascertaining whether the 99% posterior probability estimates of the α and β parameters included zero (i.e., by identifying positive or negative non-zero estimates of the parameters). In addition, loci which were extreme ‘introgression outliers’ were identified for both parameters by identifying loci whose median estimates were not included in the 99% posterior probability credible intervals (Gompert & Buerkle 2011). Further details of visualising shared variation, hybrid category assignment and *bgc* analysis are shown in Supporting Information Methods S2.

## Results

### Phylogenetic analysis

Using DNA sequence data from 226 nuclear gene regions, we produced a ‘species’ tree using *ASTRAL* (Zhang *et al*., 2018) (Fig. 2) based on gene trees inferred with the program *RAxML* (Stamatakis 2014). This resulted in well-supported relationships between major subclades (>0.9 local posterior probability (PP)), with lower support for inter-specific relationships. *ASTRAL* inferred a high degree of discordance among gene tree topologies at many nodes (quartet score = 0.52), with multiple alternative bipartitions reconstructed at most nodes. This is evident from the presence of many more conflicting gene trees (numbers below branches) than congruent gene trees (numbers above branches) in Supporting Information Fig. S1.

**Figure 2:**
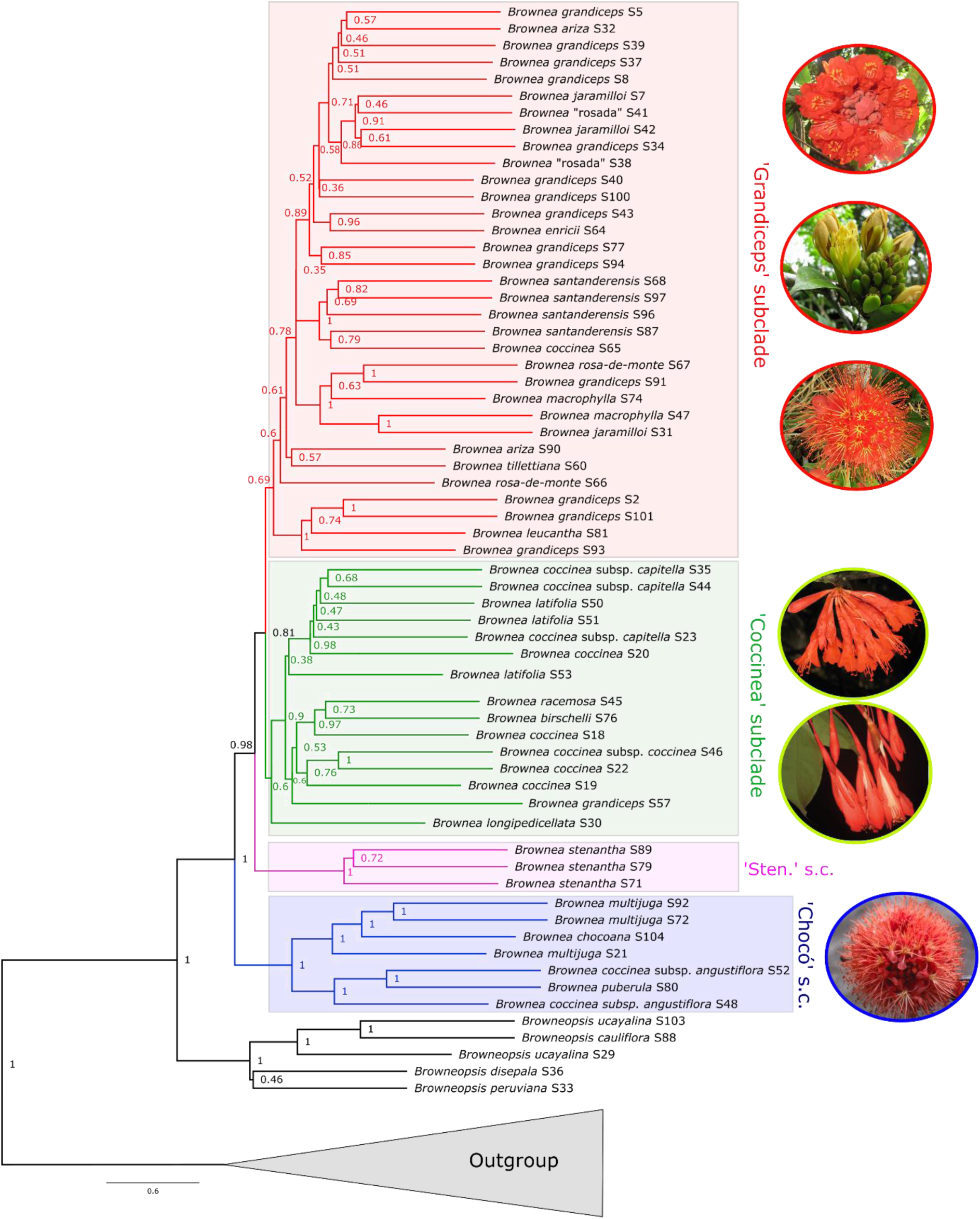
*ASTRAL* species tree inferred from *RAxML* gene trees using the multi-species coalescent model. Numbers at nodes of the tree signify local posterior probability (LPP) values, a measure of support for each quadripartition. Taxa within the red box belong to the ‘Grandiceps’ subclade, taxa within the green box belong to the ‘Coccinea’ subclade, taxa within the pink box represent the ‘Stenantha’ subclade and those within the blue box belong to the ‘Chocóan’ subclade. Images show inflorescences of species within *Brownea*-species shown are, from top to bottom: *B. grandiceps* (photograph © Rowan Schley), *B. jaramilloi* (photograph © Xavier Cornejo), *B. macrophylla* (photograph © Bente Klitgaard), *B. coccinea* subsp. *capitella* (photograph © Xavier Cornejo), *B. longipedicellata* (photograph © Domingos Cardoso) and *B. multijuga* (photograph © Bente Klitgaard).

### Ancient introgression in Brownea species

Phylogenetic networks estimated to ascertain whether diversification in *Brownea* was tree-like or reticulate pinpointed two hybridization events within the genus (Fig. 3). This is indicated by the −log pseudolikelihood values in Supporting Information Fig. S2, since the number of hybridization events (*h*) which best describe the data give the largest improvement in −log pseudolikelihood. In Fig. S2, − log pseudolikelihoods increased steadily between *h* = 0 and *h* = 2, after which the increasing number of hybridization events only made minimal improvements to −log pseudolikelihood.

**Figure 3:**
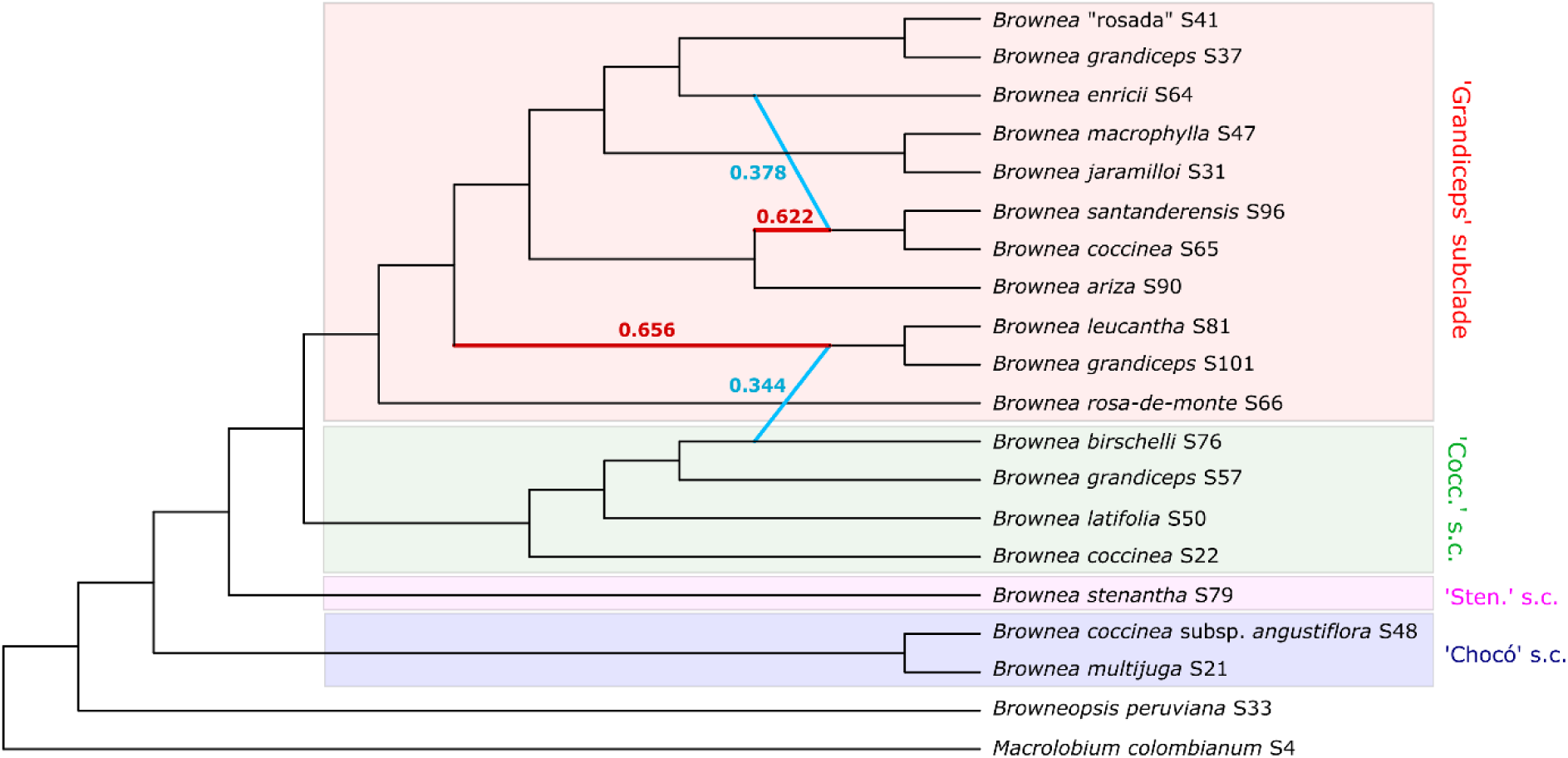
Phylogenetic network with two hybridization events (*h* = 2), estimated using *SNaQ!* in the Julia package *PhyloNetworks*. Light blue horizontal branches indicate inferred hybridization events, and numbers next to the branches show the estimated proportion of genes contributed by each lineage in the hybridization event. Red branches signify the ancestral lineage and what proportion of the modern lineage’s genes were contributed from it. Taxa were chosen in order to represent one accession per lineage, as per the assumptions of *PhyloNetworks*, and the individual with the highest gene recovery was used for each lineage. Coloured boxes containing taxa are described in the legend of Fig. 2.

The inferred phylogenetic network (Fig. 3) showed a broadly similar topology to the species tree in Fig. 2, with the addition of two hybridization events. The first hybridization event occurred between the lineage leading to the Venezuelan accessions of *B. grandiceps/B. leucantha* and *B. birschelli*, suggesting that the lineage leading to *B. birschelli* has in the past contributed around 34% of the genetic material present in the common ancestor of the Venezuelan *B. grandiceps* and *B. leucantha*. The second inferred occurrence of hybrid ancestry occurs between the ancestors of the subclade containing the Colombian accessions of *B. coccinea/B. santanderensis* and the lineage leading to *B. enricii*, which contributed around 37% of the genetic material belonging to the ancestor of the aforementioned subclade.

Tree Incongruence Checking in R (TICR) analysis indicated that a network-like model (i.e., one including introgression) best described the patterns of incongruence in the single-accession-per-species gene trees inferred during this study. This method suggested an excess of outlier quartets (*P=* 1.240 × 10^−19^, *Χ*^2^ = 91.149), which differed significantly from the CF values expected under a coalescent model describing only incomplete lineage sorting. As such, a tree-like model was able to be rejected as an explanation for the observed relationships between taxa in *Brownea*, suggesting that hybridization occurred over the course of its evolutionary history and was responsible for the observed incongruence.

### Introgression dynamics across loci

Our population genomic analyses revealed a broad degree of shared variation between *B. grandiceps* and *B. jaramilloi*, along with a homogeneous distribution of introgression rates among loci. In total, ddRAD sequencing resulted in 640,898,950 reads between 350-550bp in length for 171 individuals. Among these reads, 28,503,796 (4.45%) were discarded due to poor recovery. There was a mean of 3,747,947 reads per individual, with an average coverage depth across all samples of 27.5x. Relatively high levels of shared genetic variation were observed among taxa, along with low levels of genetic differentiation and only marginal differences in the amount of genetic variation, as shown by the population genetic statistics calculated by the Stacks pipeline for the two *Brownea* species (including *B*. “rosada”, which is grouped with *B. grandiceps*) (Supporting Information Table S3). The F_st_ calculated between *B. grandiceps/B*. “rosada” and *B. jaramilloi* was 0.111, representing a low degree of fixation, and so a high amount of shared variation.

Patterns of shared genetic variation visualised with principal component analysis (PCA) and *SPLITSTREE* showed a distinct signature of admixture between *B. grandiceps* and *B. jaramilloi*. The PCA of SNP variation inferred using the R package *ggplot2* (Fig. 4a) indicated that the first principal component (PC1) explained the largest proportion of the genetic variation among all the principal components (20.4%). Individuals of *B. grandiceps* are tightly clustered along PC1, where the individuals of *B. jaramilloi* show much more variability, with many individuals forming an intergradation between the two main species clusters. Additionally, the *B*. “rosada” accessions appear to have clustered more closely to *B. grandiceps* than to *B. jaramilloi* along PC1. PC2, which explained 10.7% of the variation in the SNP data, shows a similar degree of variability in both *B. grandiceps* and *B. jaramilloi*, with two accessions of *B*. “rosada” shown to cluster in between both species. This pattern is reflected by the implicit network built using *SPLITSTREE* (Supporting Information Fig. S3). *SPLITSTREE* recovered a clustering of individuals into two main groups, largely representing *B. grandiceps* and *B. jaramilloi*, with four putative hybrid individuals displaying an intermediate relationship between the two species clusters.

**Figure 4:**
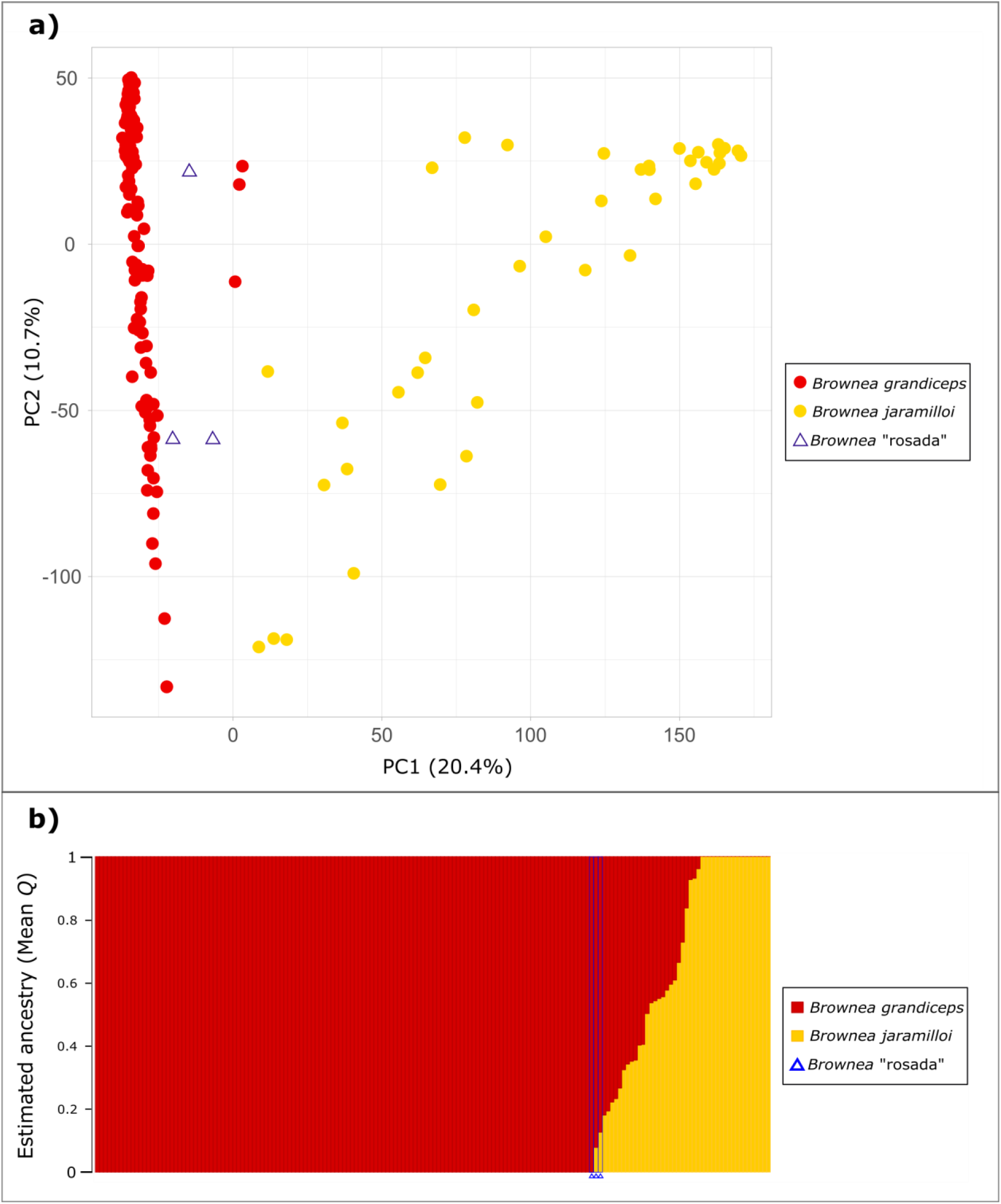
**a)** Principal component analysis of genotype data for all SNPs inferred at all loci. Individuals are coded by colour and point shape: red circles denote individuals identified as *B. grandiceps*, yellow circles denote individuals identified as *B. jaramilloi*, and blue triangles denote putative hybrid individuals (*B*. “rosada”). The amount of the genetic variation explained by each principle component is shown next to axes, which are labelled PC1 and PC2. **b)** *fastSTRUCTURE* plot, indicating ancestry proportions for two populations (*K* = 2). The ‘populations’ correspond to two different species: *B. grandiceps* (shown in red) and *B. jaramilloi* (shown in yellow). Each individual accession is represented by a column, and the proportion of ancestry from either species (mean *Q*) is proportional to the length of different coloured bars in each column. Individuals identified as *B*. “rosada” are marked with a blue box and a blue triangle beneath their column. All taxa to the left and right of these boxes were identified as *B. grandiceps* and *B. jaramilloi*, respectively.

The *fastSTRUCTURE* analysis (Raj *et al*., 2014) performed to further examine the degree of shared genetic variation indicated that there was a large amount of shared ancestry between the genotyped individuals (Fig. 4b), with evidence of extensive backcrossing due to the widely differing proportions of ancestry in different individuals. Marginal likelihood comparison indicated that the best value of *K* (i.e., the number of populations) was two (Δ_marginal likelihood_ = 0.085, Supporting Information Fig. S4A), which was the value that resulted in the largest increase in marginal likelihood. Since the ‘best’ value of *K* is an estimate, f*astSTRUCTURE* plots generated with other *K* values are shown in Supporting Information Fig. S4B. Figure 4b shows that most individuals of *B. jaramilloi* have varying fractions of *B. grandiceps* ancestry in their genome. In addition, the individuals identified as *B*. “rosada” appear to have inherited most of their ancestry from *B. grandiceps*, with only a minimal contribution from *B. jaramilloi*. The same pattern was recovered from a *fastSTRUCTURE* run incorporating 40 random individuals from each species, performed to account for any bias which may have been incurred by differences in sample size (Supporting Information Fig. S4C).

The *NEWHYBRIDS* analysis revealed that most hybrid individuals were the result of continued hybridization, with pure *B. grandiceps* making up 73.9% of the genotyped population, and pure *B. jaramilloi* making up 21.9% for the 500 subsampled loci used in this analysis. There were no F1 (first generation) hybrids identified by the subset of loci analysed, and all hybrid individuals were either F2 hybrids (0.592%) or had a broad distribution of probabilities across hybrid classes (3.55%), suggesting backcrossing. In the latter, the probability of any one hybrid class did not exceed 90%, which was the threshold used to categorize individuals as belonging to a certain class.

The Bayesian estimation of genomic clines (*bgc*) analysis, which was used to determine whether any loci showed outlying rates of admixture relative to the genome-wide average, recovered a signal of asymmetric introgression, but did not detect any differential rates of introgression between loci. Of the 19,130 loci under study, *bgc* recovered 251 loci (1.3%) with mostly *B. jaramilloi* alleles among all individuals across species (i.e. with positive α estimates) and 1089 loci (5.69%) with mostly *B. grandiceps* alleles among all individuals across species (i.e., with negative α estimates). However, no loci displayed extreme rates of introgression relative to the average rate across the genome (i.e., there were no statistically outlying β parameter estimates). The MCMC runs from which these results were drawn showed adequate mixing (Supporting Information Fig. S5A-C).

## Discussion

Our study represents the first clear case of reticulated evolution among Amazonian trees documented at multiple phylogenetic levels. This contrasts with previous work, which suggested that hybridization is an extremely rare phenomenon in Amazonian trees. We demonstrate that within *Brownea*, a characteristic Amazonian tree genus, reticulated evolution has occurred over the course of its evolutionary history, with evidence of introgression deep in the history of the genus and more recently, between the closely related *B. grandiceps* and *B. jaramilloi*.

### *Introgression has occurred at deep phylogenetic levels in* Brownea

Phylogenetic network analysis suggested that introgression has taken place between *Brownea* lineages in the past, with two separate hybridization events inferred (Fig. 3). The concordance factors obtained from gene trees were not adequately explained by a tree-like model (i.e., one accounting only for incomplete lineage sorting), suggesting that a network-like model (i.e., one including introgression) best describes the diversification patterns within *Brownea* (*P=* 1.240 × 10^−19^, *Χ*^2^ = 91.149). Hybridization mainly occurs between individuals which are geographically overlapping, or have been so at some point in the past (Burbrink & Gehara 2018). Our findings are thus corroborated by the fact that both historical introgression events were between taxa that co-occur (e.g., *B. grandiceps, B. leucantha* and *B. birschelli* on the Northern coast of Venezuela, and *B. coccinea, B. santanderensis* and *B. enricii* within the Colombian cordilleras). This is also in agreement with previous work (Schley *et al*., 2018), which indicated that the ‘stem’ lineages within *Brownea* shared a broad distribution across northern South America, which may have further facilitated earlier hybridization. Our study also recovered a low degree of support for bifurcating inter-specific relationships (Fig. 2) and a high degree of incongruence among gene trees (Supporting Information Fig. S1). These results can be partially explained by introgression, rather than being caused purely by the stochasticity of lineage sorting in large populations, which is suggested to be a common phenomenon in many rainforest trees with large population sizes (Pennington & Lavin 2016). It seems likely that amongst closely-related species of *Brownea* divergence has not progressed to the point of complete reproductive isolation, as has been shown in temperate *Quercus* species (e.g. Lepais & Gerber 2011). Moreover, the low quartet scores and minimal gene-tree concordance found at the species level in this study mirror those observed in *Lachemilla*, a montane neotropical plant genus, in which gene tree discordance was shown to be explained by both historical and recent hybridization (Morales-Briones, Liston, & Tank 2018).

Since the inferred events of ancient introgression both involved the common ancestors of two sets of present-day species, they are likely to have occurred several million years in the past, before the divergence of the descendent species. The inferred introgression events are likely to have occurred since the Miocene period (∼23Ma), as date estimates from previous work suggest that *Brownea* originated around this time (Schley *et al*., 2018). Accordingly, it is unlikely that the inferred hybridization events are due to ‘accidental’ hybridization between co-occurring species, merely producing maladapted F1 hybrids which will eventually be outcompeted, fortifying the boundaries between species (i.e. ‘reinforcement’ (Hopkins 2013)). Rather, our results suggest a persistence of introgression over evolutionary time within *Brownea*.

### *Recent hybridization also occurs between* Brownea *species, resulting in persistent hybrids and introgression across the genome*

This investigation also uncovered a substantial signal of recent introgression between *B. grandiceps* and *B. jaramilloi* in the Yasuní National Park 50ha plot located in the Ecuadorian Amazon. The low F_st_ estimate for the *B. grandiceps/B*. “rosada” and *B. jaramilloi* populations (0.11), in addition to the principal component analysis (Fig. 4a), the *fastSTRUCTURE* analysis (Fig. 4b) and the *SPLITSTREE* plot (Supporting Information Fig. S3) indicated a high degree of shared variation between these lineages. All these approaches show that individuals cluster into two main groups, largely representing *B. grandiceps* and *B. jaramilloi*, as well as a set of individuals forming an intergradation between the two main clusters. The individuals which form a part of this intergradation were mostly identified as *B. jaramilloi*, although hybrid individuals are also observed in the *B. grandiceps* cluster. The higher number of variant sites and the higher nucleotide diversity (π) recovered for *B. jaramilloi* (Supporting Information Table S3) could also reflect the hybrid ancestry of many individuals identified as this species. A similar introgression-driven increase in nucleotide diversity has been shown in closely-related species of *Mimulus* which undergo asymmetric introgression (Sweigart & Willis 2003). In addition, *B. jaramilloi* and *B. grandiceps* showed evidence of multi-generational backcrossing in both the *fastSTRUCTURE* and *NEWHYBRIDS* analyses, further suggesting that hybridization does not always result in reinforcement and that hybrids persist over time.

The *bgc* analysis indicated that most introgression was asymmetric, with more loci containing mostly *B. grandiceps* alleles (5.96% of all loci) than loci containing mostly *B. jaramilloi* alleles (1.3% of all loci). Importantly, this *bgc* analysis also indicated that gene flow occurs largely at the same rate across loci, since there were no outlying estimates of the β parameter. The excess of *B. grandiceps* ancestry mirrors the asymmetry of introgression suggested by Fig. 4b and is likely driven by the uneven population sizes of the two species. *Brownea jaramilloi* has only ever been observed in a small part of the western Amazon (Pérez *et al*., 2013) and is much less populous than *B. grandiceps*, which occurs across northern South America. More specifically, the disproportionate donation of alleles from *B. grandiceps* may be due to ‘pollen swamping’, whereby pollen transfer from another, more populous species is more likely than pollen transfer from less numerous conspecifics (Buggs & Pannell 2006). Pollen swamping can serve as a mechanism of invasion (e.g., in *Quercus* (Petit *et al*., 2004)), and the associated asymmetric introgression may lead to the extinction of the rarer species, especially when hybrids are inviable or sterile (Balao *et al*., 2015). However, due to the ongoing backcrossing and introgression evident between *B. grandiceps* and *B. jaramilloi* it appears that hybrids are not always sterile, or at least that ‘foreign’ alleles are not always subject to strong purifying selection. The best evidence of this is the absence of loci with extreme values of the β parameter recovered by our *bgc* analysis. Extreme deviations in β are mainly expected in the presence of gene flow when selection against hybrids is strong, resulting in underdominance and a paucity of heterozygous sites (Gompert *et al*., 2012). As such, our inferred lack of outlying β values suggests persistence of hybrid genotypes.

Further to this, the observed asymmetry in introgression could be caused by selection favouring hybrid genotypes. The transfer of genomic regions which confer a selective advantage from a donor to a recipient species (i.e. adaptive introgression) can result in a disproportionate amount of one species’ genome being present in hybrid individuals (Yang *et al*., 2018). This occurs when viable hybrids only tend to backcross with one of the parental species, thereby introducing the other species’ genetic material into the first in an asymmetrical fashion. Selection has been suggested as a driver for asymmetric introgression in multiple plant genera, including *Helianthus* (Scascitelli *et al*., 2010) and *Iris* (Arnold *et al*., 2010). Accordingly, it is possible that selectively advantageous alleles are passing between species (e.g., between *B. jaramilloi* and *B. grandiceps*) through backcrossing events over many generations as observed by the *fastSTRUCTURE* and *NEWHYBRIDS* analyses in this study (Mallet 2005). This may facilitate adaptation to new niches due to the widening of the pool of variation upon which selection can act (e.g. Whitney *et al*., 2010). However, it is difficult to ascertain whether adaptive introgression has occurred without measuring the impact of introgression on variation in the phenotype and its fitness effects (Suarez-Gonzalez *et al*., 2018).

### The contribution of hybridization to neotropical rainforest tree diversity

Studies such as ours that document persistent hybridization across evolutionary time between tree species within Amazonia, and within tropical rainforests in general, are rare. Indeed, it was previously suggested that inter-specific hybrids between rainforest tree species are poor competitors, and that fertile hybrid populations are nearly non-existent (Ashton 1969). While there is some evidence of introgression in tropical trees (Scotti-Saintagne *et al*., 2013), most available studies substantiating this are based on trees from other tropical regions or habitats (e.g., *Shorea* in Asia (Kamiya *et al*., 2011; Kenzo *et al*., 2019), or *Rhizophora* in Indo-Pacific mangroves (Lo 2010)). Many of these other instances appear to occur only in degraded habitats, or involve infertile first-generation hybrids with minimal backcrossing, which contrasts with the findings of our study. Within *Brownea* we uncovered introgression across taxonomic, spatial and temporal scales, with evidence of backcrossing and the persistence of hybrid genotypes.

It is possible that a lack of selection against hybrids allows the passage of variation between *Brownea* species, which may persist over evolutionary time and could explain why we found evidence of introgression at both macroevolutionary and microevolutionary scales. This is congruent with a ‘syngameon’ model, where large, disparate populations of individuals belonging to multiple interfertile species occasionally hybridize and exchange genetic variants. This can aid in the maintenance of genetic diversity in species which exist in very low population densities, as is typical of tropical rainforest trees, thereby preventing Allee effects. This has been shown in temperate tree species (Cronk & Suarez-Gonzalez 2018), although it typically occurs at the edge of species ranges and its prevalence in hyperdiverse systems such as tropical rainforests is a topic of recent debate (e.g. Cannon & Lerdau 2019). Furthermore, while the passage of adaptive variation between syngameons of closely related species can seed rapid, speciose radiations in other taxa (Marques *et al*., 2019; Meier *et al*., 2017), whether this occurs between tree species in neotropical rainforest remains a key knowledge gap.

While it is difficult to determine whether hybridization between tree species occurs across many syngameons of closely related Amazonian lineages, or whether it is a tendency unique to certain genera such as *Brownea*, it may be prudent to review how relationships among tropical tree lineages are inferred. Accordingly, the use of phylogenetic networks in resolving the relationships between such groups may provide additional insight into whether reticulate evolution has contributed to diversification within Amazonian rainforest, which is among the most species-rich environments on Earth (Burbrink & Gehara 2018; Solís-Lemus *et al*., 2017).

## Acknowledgements

Laboratory work, fieldwork and herbarium visits to NY and US were funded by the NERC SSCP DTP, and a Genetics society Heredity Fieldwork grant funded part of the fieldwork in Ecuador. Phylogenomic sequencing was also part-funded by a grant from the Royal Botanic Gardens, Kew. Thank you to Dario Ojeda for baits and associated help, to Laszlo Csiba, Penny Malakasi, Niroshini Epitawalage, Dion Devey, Robyn Cowan and Steven Dodsworth for help in the laboratory, and to Gregor Gilfillan at the Norwegian sequencing centre for help with ddRAD sequencing. Thank you to Andres Melo Burbano for his help during fieldwork, as well as to Renato Valencia and Álvaro Pérez for helping to coordinate fieldwork in Ecuador. Thanks to Colin Hughes, Jeff Doyle and Matteo Fumagalli for comments on analytical methods, and to Peter Raven and Benjamin Torke for their comments on the rarity of hybridization in tropical trees.

Collection, transport and extraction of genetic material from plant accessions collected in Yasuní National Park were authorised by Ecuador’s Environment Ministry. Permits were issued with the following authorization numbers:

- **Collection permit:** 021-2016-IC-FAU-FLO-DPAO-PNY;
- **Mobilization permit**: 037-2016-MOV-FLO-MAE-DPAO;
- **Export permit**: 208-2019-EXP-CM-FAU-DNB/MA; and
- **Genetic research permit (Contrato Marco)**: MAE-DNB-CM-2018-0082

## Data Accessibility

The data that support the findings of this study are openly available from online repositories. All raw reads generated with the targeted bait capture and ddRADseq methods are available on the NCBI Sequence Read Archive with the accession numbers SAMN13439069-SAMN13439140 and SAMN13441804-SAMN13441974, respectively, under the Bioproject number PRJNA592723. All full phylogenomic sequence alignments, single-accession-per-species alignments and tree files, bgc input files, Stacks output files and the Detarioideae bait kit sequence file are found on Dryad (https://doi.org/10.5061/dryad.k3j9kd53w). Data are under embargo until publication, and any further data required are available from the corresponding author upon reasonable request.

## Author Contributions

R.J.S. conceived this study and performed all analyses, supervised by T.B., F.F. and B.K. Population genomic data were generated by R.J.S., and phylogenomic data were generated by R.J.S., I.K. and I.L. Baits for hybrid capture were provided by M.d.l.E. R.J.S. wrote the first draft of the manuscript with contributions from R.T.P., O.A.P.E., A.J.H., M.d.l.E., I.L., T.B., F.F. and B.K.

