## Supplementary Information for "Introgression across evolutionary scales suggests reticulation contributes to Amazonian tree diversity"

MOLECULAR ECOLOGY

### Table of Contents:


| | |
|---|---|
| **Fig. S1** | Page 2 |
| **Fig. S2** | Page 3 |
| **Fig. S3** | Page 4 |
| **Fig. S4** | Page 5 |
| **Fig. S5** | Page 8 |
| **Table S1** | Page 10 |
| **Table S2** | Page 13 |
| **Table S3** | Page 18 |
| **Methods S1** | Page 19 |
| **Methods S2** | Page 23 |

MOLECULAR ECOLOGY

**Fig. S1** *PhyParts* analysis based on the *ASTRAL* species tree, inferred from *RAxML* gene trees using the multi-species coalescent model. Pie charts at nodes show the number of gene trees with a congruent topology at a node (blue segments), the number which support a single alternate topology (green segments), the number which support all other conflicting bipartitions (red segments) and how many gene trees were uninformative at the node (grey segments). Numbers above branches show how many gene trees were congruent at a node, whereas numbers below branches show the number of conflicting gene trees at a node.

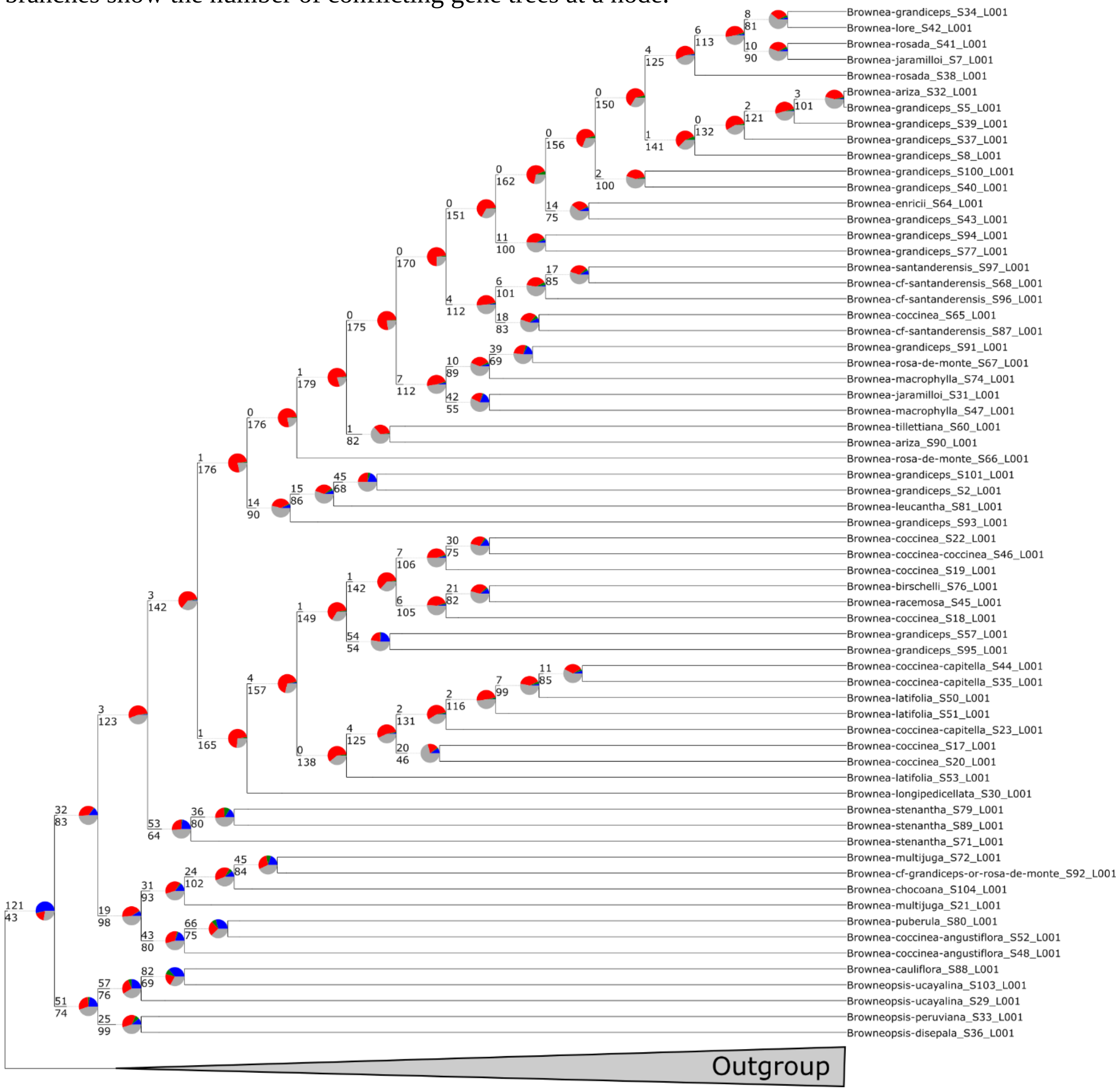

**Fig. S2** Negative log pseudolikelihood profile for each number of hybridization events inferred using *SNaQ!*. The best-fitting number of hybridization events (*h*) is displayed as the value at which the rate of change in -log pseudolikelihood plateaus, which in this case was *h=2*.

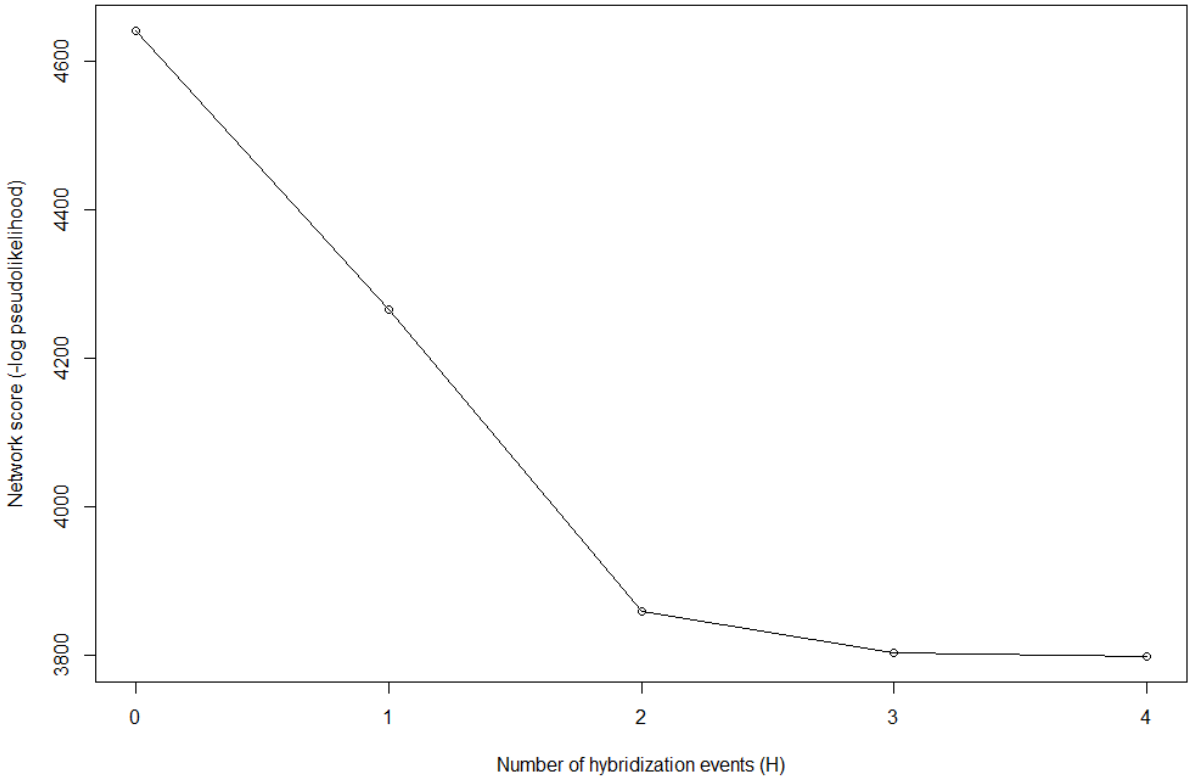

MOLECULAR ECOLOGY

**Fig. S3** *SPLITSTREE* built using uncorrelated P distances based on the full dataset for all genotyped individuals. The three coloured zones represent groupings of the two species (*B. jaramilloi* and *B. grandiceps*) as well as hybrid individuals. Groupings in the coloured zones were defined based on the clustering apparent in this plot, not on the species delimitations present in Table S4.

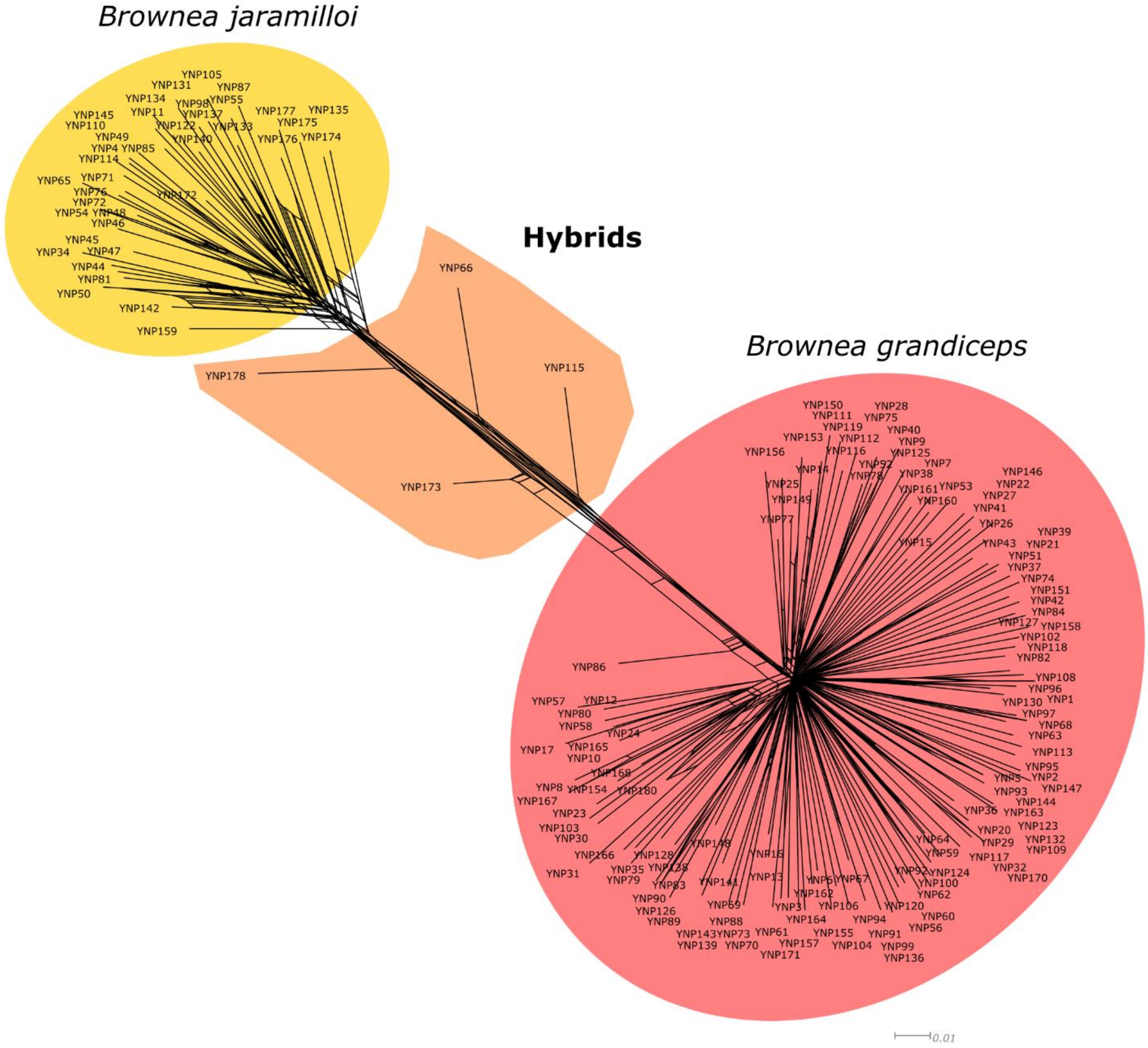

**Fig. S4 A)** Difference in marginal likelihoods between sequential numbers of population clusters (*K*), as estimated by *FastSTRUCTURE.* The best value of *K* results in the largest increase in marginal likelihood (in this case, *K=2*).
**B)** *fastSTRUCTURE* plots generated using different values of estimated clusters (*K*=3 to *K*=5). This plot indicates ancestry proportions from *K* inferred population clusters. Each individual accession is represented by a column, and the proportion of ancestry from each inferred population is proportional to the length of different coloured bars in each column.
**C)** *fastSTRUCTURE* plot incorporating 40 individuals from each of the two species under study in order to reduce error associated with different sample sizes. This plot indicates ancestry proportions from *K* = 2 population clusters (which are, in this case, different species: *B. grandiceps* and *B. jaramilloi*).  Each individual accession is represented by a column, and the proportion of ancestry from either species is proportional to the length of different coloured bars in each column.

**A)**

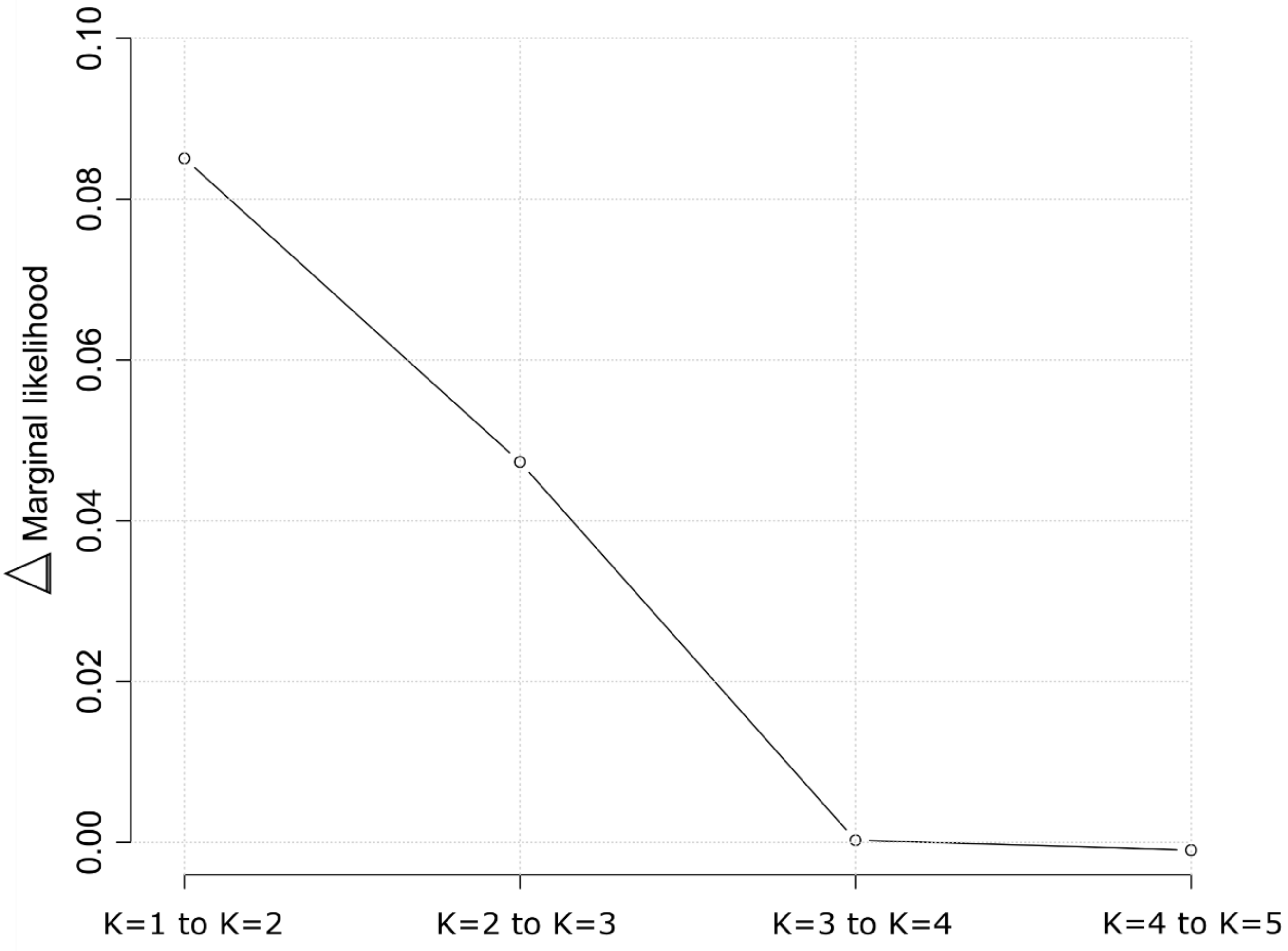

B)

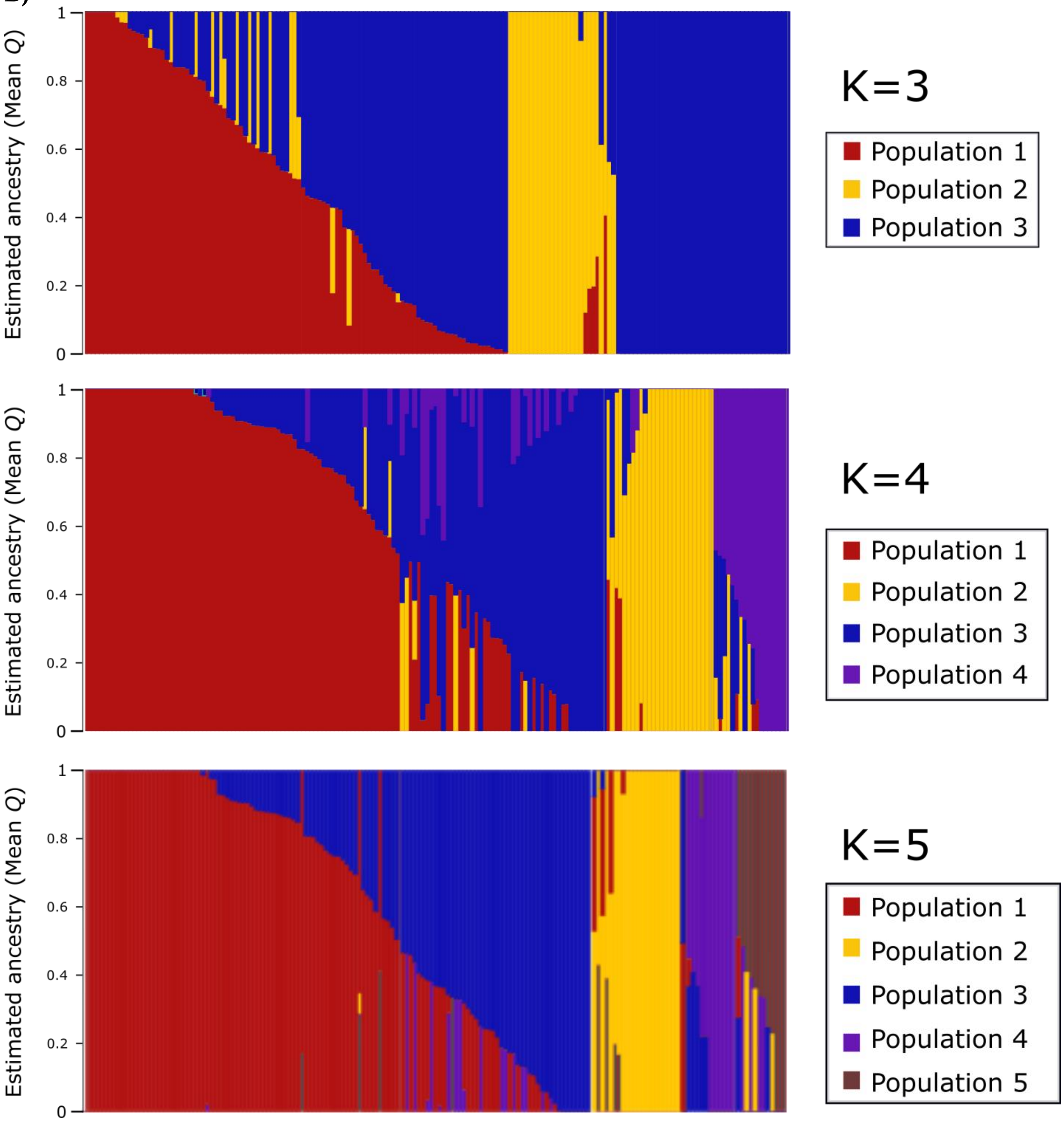
Estimated ancestry (Mean Q)
1
0.8
0.6
0.4
0.2
0
K=3
Population 1
Population 2
Population 3
K=4
Population 1
Population 2
Population 3
Population 4
K=5
Population 1
Population 2
Population 3
Population 4
Population 5

C)

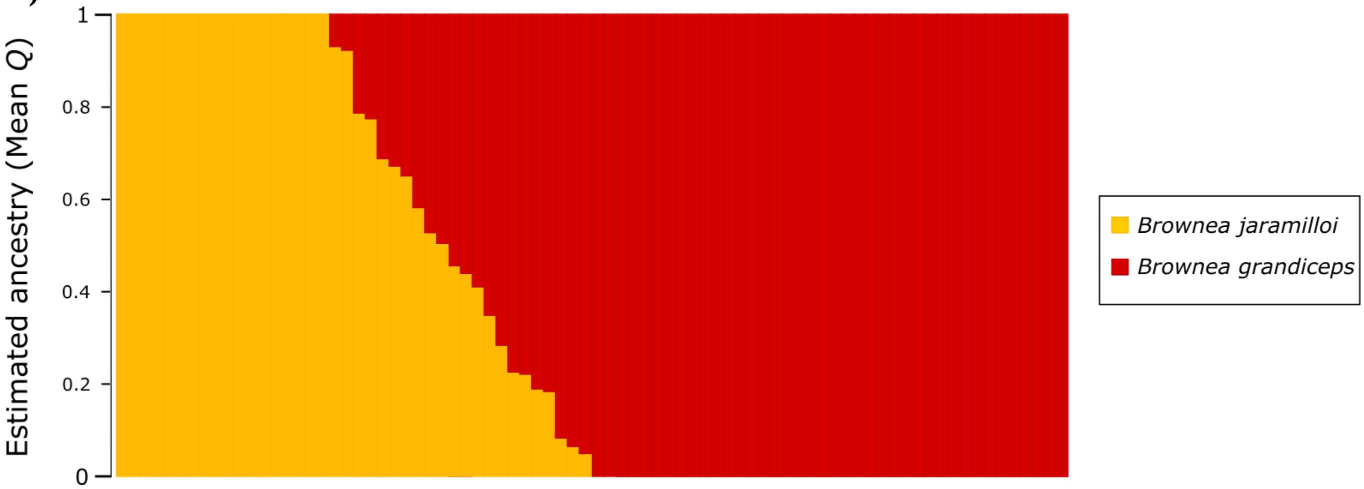
Estimated ancestry (Mean Q)
1
0.8
0.6
0.4
0.2
0
Brownea jaramilloi
Brownea grandiceps

**Fig. S5** Assessment of MCMC chain mixing for runs used to make posterior estimates for the α and β parameters in *bgc*.
**A)** Log-likelihood plot, showing log-likelihood plotted against number of MCMC iterations.
**B)** Histogram of Geweke's diagnostics for the MCMC chains used to estimate the α parameter, with the significance cut-offs for the Geweke's diagnostic ($Z$ = +/- 1.96) shown by blue dashed lines.
**C)** Histogram of Geweke's diagnostics for the MCMC chains used to estimate the β parameter, with the significance cut-offs for the Geweke's diagnostic ($Z$ = +/- 1.96) shown by blue dashed lines.

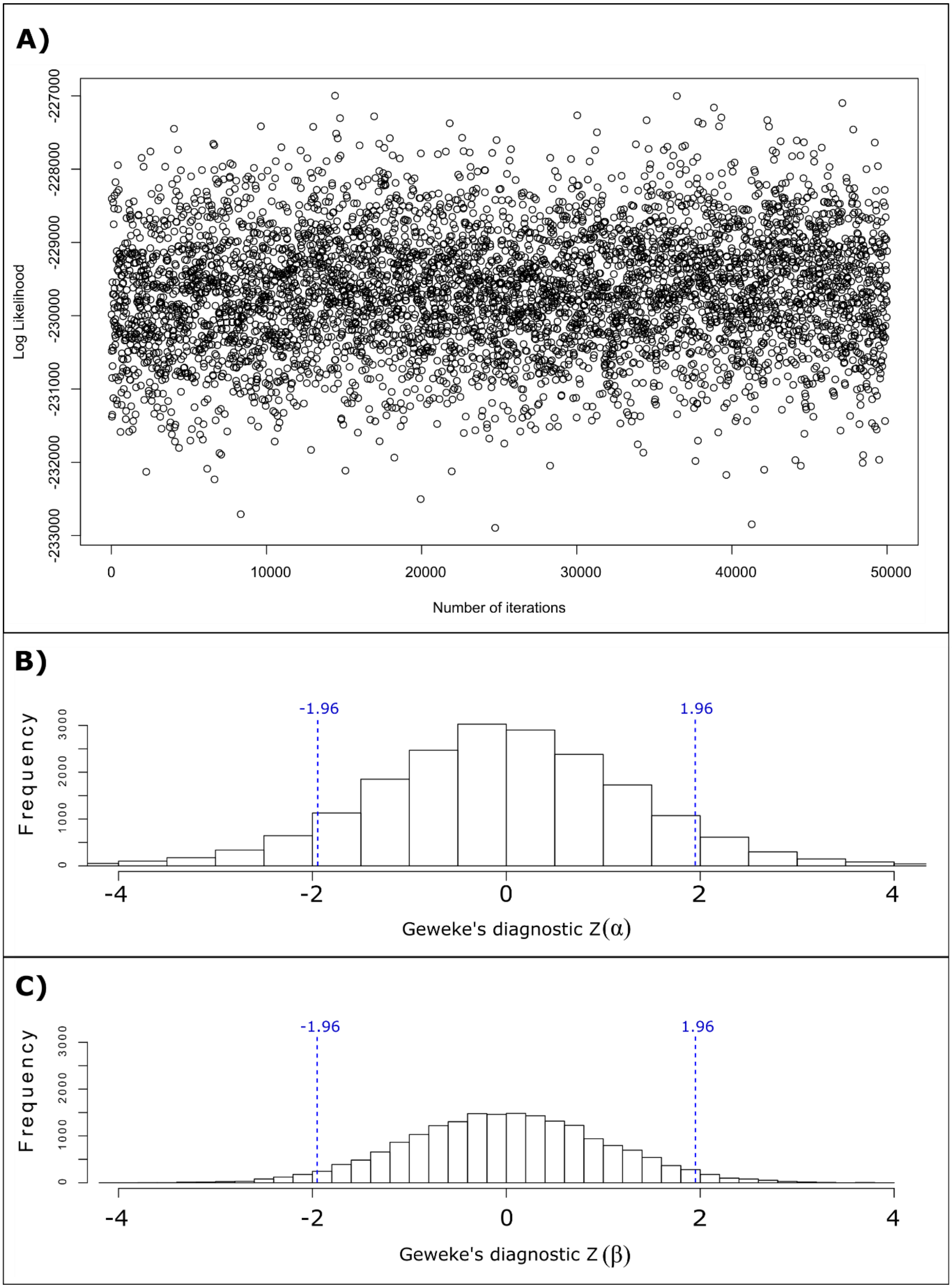
A)
Log Likelihood
-227000
-228000
-229000
-230000
-231000
-232000
-233000
0
10000
20000
30000
40000
50000
Number of iterations
B)
Frequency
0
1000
2000
3000
-1.96
1.96
-4
-2
0
2
4
Geweke's diagnostic Z(α)
C)
Frequency
0
1000
2000
3000
-1.96
1.96
-4
-2
0
2
4
Geweke's diagnostic Z (β)

MOLECULAR ECOLOGY

**Table S1** DNA numbers, species identifications, sampled vouchers, collector information and collection localities for *Brownea* species (and Brownea clade outgroup species) used in this study. Herbaria from which accessions were collected are cited after the collector name and number. Outgroup taxa are marked with an asterisk (*).

| DNA Number | Identification | Voucher | Collection locality |
|---|---|---|---|
| S1 | *Macrolobium discolor* Benth.* | Iganci 886 (MO) | Brazil |
| S2 | *Brownea grandiceps* Jacq. | Edwards 328 (K) | Aragua, Venezuela |
| S3 | *Macrolobium montanum* var. *potaroanum* R.S. Cowan* | Andel 5548 (K) | Suriname |
| S4 | *Macrolobium colombianum* (Britton & Killip) Killip ex L. Uribe* | Klitgaard 682 (K) | Napo, Ecuador |
| S5 | *Brownea grandiceps* Jacq. | Klitgaard 99504 (AAU) | Napo, Ecuador |
| S6 | *Heterostemon mimosoides* Desf.* | Iganci 875 (MO) | Brazil |
| S7 | *Brownea jaramilloi* A.J. Pérez & Klitg. | Klitgaard 662 (K) | Yasuní, Napo, Ecuador |
| S8 | *Brownea grandiceps* Jacq. | Daly 6167 (NY) | Rio Marañon, Loreto, Peru |
| S18 | *Brownea coccinea* Jacq. | Edwards 372 (K) | Aragua, Venezuela |
| S19 | *Brownea coccinea* subsp. *coccinea* Jacq. | Steyermark 126503 (K) | Morrocoy, Venezuela |
| S20 | *Brownea coccinea* subsp. *capitella* (Jacq.) D. Velazquez | Clark 6062 (K) | Cuyuni-Mazaruni, Guyana |
| S21 | *Brownea multijuga* Britton & Killip | Pennington 15642 (K) | Pichincha, Ecuador |
| S22 | *Brownea coccinea* Jacq. | Trujillo 19314 (K) | Falcon, Venezuela |
| S23 | *Brownea coccinea* subsp. *capitella* (Jacq.) D. Velazquez | Gutierrez (K) | Venezuela |
| S24 | *Macrolobium herrerae* Zarucchi* | Marshall 392 (K) | Izabal, Guatemala |
| S25 | *Macrolobium parvifolium* (Huber) R.S. Cowan* | de Lima 6814 (K) | Acre, Brazil |
| S26 | *Macrolobium longipedicellatum* Ducke* | de Lima 2790 (K) | Amazonas, Brazil |
| S27 | *Paloue speciosa* (Ducke) Redden* | Martins 54 (K) | Amazonas, Brazil |
| S28 | *Paloue princeps* (Schomb. Ex Benth.) Redden* | Sabatier (K) | French Guiana |
| S29 | *Browneopsis ucayalina* Huber* | Klitgaard 684 (K) | Morona-Santiago, Ecuador |
| S30 | *Brownea longipedicellata* Huber | Klitgaard (Silica) | Rio de Janeiro Botanical Garden, Brazil |

| | | | |
|---|---|---|---|
| S31 | *Brownea jaramilloi* A.J. Pérez & Klitg. | Villa 1645 (MO) | Yasuní, Napo, Ecuador |
| S32 | *Brownea ariza* Benth. | Valencia 68473 (AAU) | Sucumbios, Ecuador |
| S33 | *Browneopsis peruviana* (J.F. Macbr.) Klitg.* | Castro MHT-01_2 (Silica) | Peru |
| S34 | *Brownea grandiceps* Jacq. | Baker HD1402 (Silica) | Ecuador |
| S35 | *Brownea coccinea* subsp. *capitella* (Jacq.) D. Velazquez | Hollowell 485 (US) | Hossororo, Barima-Waini, Guyana |
| S36 | *Browneopsis disepala* (Little) Klitg.* | Klitgaard 67032 (AAU) | Rio Palenque, Los Rios, Ecuador |
| S37 | *Brownea grandiceps* Jacq. | Villa 1855 (Silica) | Yasuní, Napo, Ecuador |
| S38 | *Brownea* "rosada" Jacq. | Villa 747 (MO) | Yasuní, Napo, Ecuador |
| S39 | *Brownea grandiceps* Jacq. | Villa 1866 (Silica) | Yasuní, Napo, Ecuador |
| S40 | *Brownea grandiceps* Jacq. | Villa 1859 (Silica) | Yasuní, Napo, Ecuador |
| S41 | *Brownea* "rosada" Jacq. | Villa 1868 (Silica) | Yasuní, Napo, Ecuador |
| S42 | *Brownea jaramilloi* A.J. Pérez & Klitg. | Villa 1609 (MO) | Yasuní, Napo, Ecuador |
| S43 | *Brownea grandiceps* Jacq. | Killip 34351 (MO) | Meta, Colombia |
| S44 | *Brownea coccinea* subsp. *capitella* (Jacq.) D. Velazquez | Maguire 46987 (NY) | El Foco, Bolivar, Venezuela |
| S45 | *Brownea coccinea* subsp. *capitella* (Jacq.) D. Velazquez | Gentry 14773 (NY) | Los Caracas, Venezuela |
| S46 | *Brownea coccinea* subsp. *coccinea* Jacq. | Gonzalez 29 (US) | Churuguara-coro, Venezuela |
| S47 | *Brownea macrophylla* hort. Ex Mast. | Villa 2004 (K) | Joya de los Sachas, Succumbios, Ecuador |
| S48 | *Brownea coccinea* subsp. *angustiflora* (Little) Klitg. | Klitgaard 99488 (AAU) | Esmeraldas, Ecuador |
| S50 | *Brownea latifolia* Jacq. | Polak 44 (K) | Guyana |
| S51 | *Brownea latifolia* Jacq. | Hoffman 2656 (US) | Pomeroon-Supenaam, Guyana |
| S52 | *Brownea coccinea* subsp. *angustiflora* (Little) Klitg. | Klitgaard 67044 (AAU) | Esmeraldas, Ecuador |
| S53 | *Brownea latifolia* Jacq. | Philcox 8078 (K) | Trinidad |
| S57 | *Brownea grandiceps* Jacq. | Dorr 7792 (NY) | Barinas, Venezuela |
| S60 | *Brownea tillettiana* D. Velásquez & G. Agostini | Davidse 18640 (NY) | Perija, Venezuela |
| S64 | *Brownea enricii* L.M. Quiñones | Garcia-Barriga 17222 (US) | Boyaca, Colombia |
| S65 | *Brownea coccinea* Jacq. | Gentry 15470 (NY) | Cimitarra, Santander, Colombia |

| | | | |
|---|---|---|---|
| S66 | *Brownea rosa-de-monte* P.J. Bergius | Callejas 3209 (NY) | Antioquia, Colombia |
| S67 | *Brownea rosa-de-monte* P.J. Bergius | Duke 11126 (3) (NY) | Rio Truando, Choco, Colombia |
| S68 | *Brownea santanderensis* L.M. Quiñones | Callejas 4503 (NY) | Antioquia, Colombia |
| S71 | *Brownea stenantha* Britton & Killip | Stern 180 (US) | Paya, Darien, Panama |
| S72 | *Brownea multijuga* Britton & Killip | Berry 5413 (NY) | Darien, Panama |
| S74 | *Brownea macrophylla* hort. ex. Mast. | Klitgaard 1802 (K) | Boyaca, Colombia |
| S76 | *Brownea birschelli* Hook. F. | Birschel *s.n.* (K) | Rio de La Guaira, Venezuela |
| S77 | *Brownea grandiceps* Jacq. | Schunke 0011 (USM 14269) (K) | Maynas, Peru |
| S79 | *Brownea stenantha* Britton & Killip | Pennell 4698 (K) | Rio Sinu, Bolivar, Colombia |
| S80 | *Brownea puberula* Little | Little 6281 (K) | Esmeraldas, Ecuador |
| S81 | *Brownea leucantha* Jacq. | Killip 37047 (K) | Miranda, Venezuela |
| S87 | *Brownea* cf. *santanderensis* L.M. Quiñones | Klitgaard 1800 (K) | Boyaca, Colombia |
| S88 | *Browneopsis cauliflora* (Poepp.) Huber* | Klug 4163 (K) | San Martin, Peru |
| S89 | *Brownea stenantha* Britton & Killip | Haught 4788 (K) | Antioquia, Colombia |
| S90 | *Brownea ariza* Benth. | Hanbury-Tracy 509 (K) | Sierra de Santa Marta, Magdalena, Colombia |
| S91 | *Brownea grandiceps* Jacq. | Allen 931 (K) | Panama province, Panama |
| S92 | *Brownea multijuga* Britton & Killip | Dawe 883 (K) | Darien, Colombia |
| S93 | *Brownea grandiceps* Jacq. | Fendler 340 (K) | Colonia Tovar, Aragua, Venezuela |
| S94 | *Brownea grandiceps* Jacq. | Grandez 1625 (K) | Loreto, Peru |
| S96 | *Brownea santanderensis* L.M. Quiñones | Fonnegra 8088 (K) | Santander, Colombia |
| S97 | *Brownea santanderensis* L.M. Quiñones | Castellanos 448 (K) | Santander, Colombia |
| S100 | *Brownea grandiceps* Jacq. | Whitmore 885 (K) | Napo/Pastaza, Ecuador |
| S101 | *Brownea grandiceps* Jacq. | Edwards 327 (K) | Aragua, Venezuela |
| S103 | *Browneopsis ucayalina* Huber* | Diaz 4228 (K) | Amazonas, Peru |
| S104 | *Brownea chocoana* (L.M. Quiñones) | Fuchs 22384 (K) | Choco, Colombia |

MOLECULAR ECOLOGY

**Table S2** DNA numbers, species, coordinates for subplots, tree tag and habitat type for individuals of *B. grandiceps*, *B. jaramilloi* and *B.* "rosada" collected from the 50-hectare forest plot in Yasuní National Park, Ecuador. The 'Subplot X' and 'Subplot Y' columns refer to the coordinates of each subplot, relative to the southwest corner of the Yasuní plot, within which each specimen was collected. 'Tag' refers to the unique identifier given to each tree with a diameter-at-breast-height of >1cm in the Yasuní plot. 'Habitat' refers to the habitat type of each subplot from which each individual was collected, with 'Valley' subplots defined as having a mean elevation ≤227.2 m, and 'ridge' subplots as having a mean elevation >227.2 m (Valencia *et al.*, 2004). Individuals marked with an asterisk (*) were from silica collections made in the Yasuní National Park 50ha plot in 2002.

| DNA number | Species | Subplot X | Subplot Y | Tag | Habitat |
|---|---|---|---|---|---|
| YNP1 | *Brownea grandiceps* Jacq. | 8.75 | 14.75 | 83337 | Ridge |
| YNP2 | *Brownea grandiceps* Jacq. | 9.25 | 12.25 | 92921 | Valley |
| YNP3 | *Brownea grandiceps* Jacq. | 49.75 | 15.5 | 494207 | Valley |
| YNP4 | *Brownea jaramilloi* Á.J. Pérez & Klitg. | 44.5 | 23 | 446861 | Ridge |
| YNP5 | *Brownea grandiceps* Jacq. | 9 | 12 | 92872 | Valley |
| YNP6 | *Brownea grandiceps* Jacq. | 48.75 | 15 | 484513 | Valley |
| YNP7 | *Brownea grandiceps* Jacq. | 33.75 | 23.5 | 335254 | Ridge |
| YNP8 | *Brownea grandiceps* Jacq. | 33.5 | 23.5 | - | Ridge |
| YNP9 | *Brownea grandiceps* Jacq. | 31.25 | 17.25 | 314007 | Valley |
| YNP10 | *Brownea grandiceps* Jacq. | 31.75 | 2.25 | 311102 | Valley |
| YNP11 | *Brownea jaramilloi* Á.J. Pérez & Klitg. | 17.75 | 18 | 174927 | Ridge |
| YNP12 | *Brownea grandiceps* Jacq. | 31.75 | 2.25 | 311108 | Valley |
| YNP13 | *Brownea grandiceps* Jacq. | 42 | 11.75 | 423844 | Ridge |
| YNP14 | *Brownea grandiceps* Jacq. | 42.75 | 11.25 | 423772 | Ridge |
| YNP15 | *Brownea grandiceps* Jacq. | 42.75 | 11.25 | 423763 | Ridge |
| YNP16 | *Brownea grandiceps* Jacq. | 42.25 | 11.5 | 423828 | Ridge |
| YNP17 | *Brownea grandiceps* Jacq. | 31.5 | 2.75 | 311160 | Valley |
| YNP20 | *Brownea grandiceps* Jacq. | 21.5 | 24 | - | Valley |
| YNP21 | *Brownea grandiceps* Jacq. | 21.5 | 24.75 | 215859 | Valley |
| YNP22 | *Brownea grandiceps* Jacq. | 16 | 23 | 165794 | Ridge |
| YNP23 | *Brownea grandiceps* Jacq. | 21 | 24.75 | 215912 | Valley |
| YNP24 | *Brownea grandiceps* Jacq. | 33 | 23.5 | 335373 | Ridge |
| YNP25 | *Brownea grandiceps* Jacq. | 35.25 | 14 | 350661 | Valley |
| YNP26 | *Brownea grandiceps* Jacq. | 16.25 | 23.5 | 165991 | Ridge |
| YNP27 | *Brownea grandiceps* Jacq. | 21 | 24.25 | - | Valley |
| YNP28 | *Brownea grandiceps* Jacq. | 38.5 | 19.5 | 384695 | Valley |
| YNP29 | *Brownea jaramilloi* Á.J. Pérez & Klitg. | 21.5 | 24 | - | Valley |
| YNP30 | *Brownea grandiceps* Jacq. | 16 | 23.75 | 166007 | Ridge |
| YNP31 | *Brownea grandiceps* Jacq. | 16.5 | 23.25 | 165882 | Ridge |

| | | | | | |
|---|---|---|---|---|---|
| YNP32 | *Brownea grandiceps* Jacq. | 21.5 | 24.25 | 215812 | Valley |
| YNP34 | *Brownea jaramilloi* Á.J. Pérez & Klitg. | 17 | 18 | - | Ridge |
| YNP35 | *Brownea grandiceps* Jacq. | 1 | 12.75 | 13576 | Valley |
| YNP36 | *Brownea grandiceps* Jacq. | 21.5 | 24.25 | 215811 | Valley |
| YNP37 | *Brownea grandiceps* Jacq. | 16 | 23 | 165789 | Ridge |
| YNP38 | *Brownea grandiceps* Jacq. | 31.5 | 17.5 | 314127 | Valley |
| YNP39 | *Brownea grandiceps* Jacq. | 21.25 | 24.25 | 215760 | Valley |
| YNP40 | *Brownea grandiceps* Jacq. | 38.75 | 19 | 384662 | Valley |
| YNP41 | *Brownea grandiceps* Jacq. | 21.75 | 24 | 215836 | Valley |
| YNP42 | *Brownea grandiceps* Jacq. | 21 | 24.25 | - | Valley |
| YNP43 | *Brownea grandiceps* Jacq. | 16.5 | 23.5 | 165966 | Ridge |
| YNP44 | *Brownea jaramilloi* Á.J. Pérez & Klitg. | 17.75 | 18 | 174933 | Ridge |
| YNP45 | *Brownea jaramilloi* Á.J. Pérez & Klitg. | 17.75 | 18.75 | - | Ridge |
| YNP46 | *Brownea grandiceps* Jacq. | 49 | 24 | - | Ridge |
| YNP47 | *Brownea jaramilloi* Á.J. Pérez & Klitg. | 49.25 | 24 | 496213 | Ridge |
| YNP48 | *Brownea jaramilloi* Á.J. Pérez & Klitg. | 49 | 24.5 | 496402 | Ridge |
| YNP49 | *Brownea jaramilloi* Á.J. Pérez & Klitg. | 49.25 | 24.25 | - | Ridge |
| YNP50 | *Brownea jaramilloi* Á.J. Pérez & Klitg. | 17.75 | 18.75 | - | Ridge |
| YNP51 | *Brownea grandiceps* Jacq. | 38 | 19.25 | 384594 | Valley |
| YNP52 | *Brownea grandiceps* Jacq. | 38 | 19 | - | Valley |
| YNP53 | *Brownea grandiceps* Jacq. | 16.25 | 23.5 | 165803 | Ridge |
| YNP54 | *Brownea jaramilloi* Á.J. Pérez & Klitg. | 49 | 24.75 | 496400 | Ridge |
| YNP55 | *Brownea jaramilloi* Á.J. Pérez & Klitg. | 28.75 | 15.75 | 284269 | Ridge |
| YNP56 | *Brownea grandiceps* Jacq. | 27.25 | 13.25 | 273375 | Valley |
| YNP57 | *Brownea grandiceps* Jacq. | 31.75 | 2.5 | 311142 | Valley |
| YNP58 | *Brownea grandiceps* Jacq. | 42.25 | 11 | 423727 | Ridge |
| YNP59 | *Brownea grandiceps* Jacq. | 24.25 | 13.25 | - | Ridge |
| YNP60 | *Brownea grandiceps* Jacq. | 24.25 | 13.75 | 243685 | Ridge |
| YNP61 | *Brownea grandiceps* Jacq. | 49.5 | 24 | 496250 | Ridge |
| YNP62 | *Brownea grandiceps* Jacq. | 24 | 13.25 | - | Ridge |
| YNP63 | *Brownea grandiceps* Jacq. | 12 | 15.5 | - | Ridge |
| YNP64 | *Brownea grandiceps* Jacq. | 24.5 | 13.5 | 243664 | Ridge |
| YNP65 | *Brownea jaramilloi* Á.J. Pérez & Klitg. | 49 | 24.25 | 496174 | Ridge |
| YNP66* | *Brownea* "rosada" Jacq. | 9.75 | 18.5 | 94514 | Ridge |
| YNP67 | *Brownea grandiceps* Jacq. | 27.25 | 13.5 | - | Valley |
| YNP68 | *Brownea grandiceps* Jacq. | 12 | 15.5 | 123716 | Ridge |
| YNP69 | *Brownea grandiceps* Jacq. | 23 | 10.5 | 233136 | Valley |
| YNP70 | *Brownea grandiceps* Jacq. | 23 | 10.75 | 233123 | Valley |
| YNP71 | *Brownea jaramilloi* Á.J. Pérez & Klitg. | 49.25 | 24.25 | 496181 | Ridge |
| YNP72 | *Brownea grandiceps* Jacq. | 49 | 24.75 | 496387 | Ridge |

| | | | | | |
|---|---|---|---|---|---|
| YNP73 | *Brownea grandiceps* Jacq. | 23 | 10.75 | - | Valley |
| YNP74 | *Brownea grandiceps* Jacq. | 16.75 | 23.75 | 165937 | Ridge |
| YNP75 | *Brownea grandiceps* Jacq. | 38.5 | 19.5 | 384698 | Valley |
| YNP76 | *Brownea jaramilloi* Á.J. Pérez & Klitg. | 49 | 24.25 | 496173 | Ridge |
| YNP77 | *Brownea grandiceps* Jacq. | 35 | 14.25 | 350346 | Valley |
| YNP78 | *Brownea jaramilloi* Á.J. Pérez & Klitg. | 38.75 | 19.25 | 384651 | Valley |
| YNP79 | *Brownea grandiceps* Jacq. | 27.75 | 17.75 | - | Ridge |
| YNP80 | *Brownea grandiceps* Jacq. | 31.5 | 17.25 | 314049 | Valley |
| YNP81 | *Brownea jaramilloi* Á.J. Pérez & Klitg. | 17.75 | 18.5 | 174947 | Ridge |
| YNP82 | *Brownea grandiceps* Jacq. | 24 | 13 | 243567 | Ridge |
| YNP83 | *Brownea grandiceps* Jacq. | 34 | 7.25 | 342122 | Valley |
| YNP84 | *Brownea jaramilloi* Á.J. Pérez & Klitg. | 49.75 | 15.75 | - | Valley |
| YNP85 | *Brownea jaramilloi* Á.J. Pérez & Klitg. | 28.75 | 15.25 | 284202 | Ridge |
| YNP86 | *Brownea grandiceps* Jacq. | 31.25 | 17 | - | Valley |
| YNP87 | *Brownea jaramilloi* Á.J. Pérez & Klitg. | 7.5 | 8.25 | 71967 | Ridge |
| YNP88 | *Brownea grandiceps* Jacq. | 23 | 10.5 | - | Valley |
| YNP89 | *Brownea grandiceps* Jacq. | 34.75 | 7.25 | 342211 | Valley |
| YNP90 | *Brownea grandiceps* Jacq. | 34.75 | 7.5 | 342248 | Valley |
| YNP91 | *Brownea grandiceps* Jacq. | 23.75 | 7.75 | 252703 | Ridge |
| YNP92 | *Brownea grandiceps* Jacq. | 29.5 | 12.25 | 290305 | Valley |
| YNP93 | *Brownea grandiceps* Jacq. | 10.25 | 13.25 | 102758 | Valley |
| YNP94 | *Brownea grandiceps* Jacq. | 27.5 | 13.5 | 273268 | Valley |
| YNP95 | *Brownea grandiceps* Jacq. | 9 | 13.5 | 97177 | Valley |
| YNP96 | *Brownea grandiceps* Jacq. | 8.75 | 14 | - | Ridge |
| YNP97 | *Brownea grandiceps* Jacq. | 8.75 | 14.5 | 83430 | Ridge |
| YNP98 | *Brownea jaramilloi* Á.J. Pérez & Klitg. | 28.5 | 15 | 284160 | Ridge |
| YNP99 | *Brownea grandiceps* Jacq. | 27.25 | 17.75 | 270493 | Ridge |
| YNP100 | *Brownea grandiceps* Jacq. | 24.25 | 13.5 | 243672 | Ridge |
| YNP102 | *Brownea grandiceps* Jacq. | 27.5 | 13.75 | 273344 | Valley |
| YNP103 | *Brownea grandiceps* Jacq. | 9.25 | 13.75 | - | Valley |
| YNP104 | *Brownea grandiceps* Jacq. | 46 | 22.75 | 465717 | Ridge |
| YNP105 | *Brownea jaramilloi* Á.J. Pérez & Klitg. | 28.25 | 15.5 | 284308 | Ridge |
| YNP106 | *Brownea grandiceps* Jacq. | 49.75 | 15.25 | 494185 | Valley |
| YNP108 | *Brownea grandiceps* Jacq. | 8.25 | 14.5 | 83365 | Ridge |
| YNP109 | *Brownea grandiceps* Jacq. | 10.25 | 13.25 | - | Valley |
| YNP110 | *Brownea jaramilloi* Á.J. Pérez & Klitg. | 44.75 | 23.5 | 446898 | Ridge |
| YNP111 | *Brownea grandiceps* Jacq. | 44.5 | 10.75 | 443597 | Ridge |
| YNP112 | *Brownea grandiceps* Jacq. | 44 | 10 | 443416 | Ridge |
| YNP113 | *Brownea grandiceps* Jacq. | 12 | 15.25 | 123459 | Ridge |
| YNP114 | *Brownea jaramilloi* Á.J. Pérez & Klitg. | 44.75 | 23.5 | 446900 | Ridge |

| | | | | | |
|---|---|---|---|---|---|
| YNP115 | *Brownea jaramilloi* Á.J. Pérez & Klitg. | 44 | 23 | - | Ridge |
| YNP116 | *Brownea grandiceps* Jacq. | 44 | 10.5 | 443678 | Ridge |
| YNP117 | *Brownea grandiceps* Jacq. | 27 | 13.75 | - | Valley |
| YNP118 | *Brownea grandiceps* Jacq. | 12 | 15.5 | 123705 | Ridge |
| YNP119 | *Brownea grandiceps* Jacq. | 44.25 | 10.25 | 443474 | Ridge |
| YNP120 | *Brownea grandiceps* Jacq. | 27.5 | 13.25 | 273277 | Valley |
| YNP122 | *Brownea grandiceps* Jacq. | 27.75 | 17.75 | 274472 | Ridge |
| YNP123 | *Brownea grandiceps* Jacq. | 27.75 | 17 | 274140 | Ridge |
| YNP124 | *Brownea grandiceps* Jacq. | 24.25 | 13.25 | - | Ridge |
| YNP125 | *Brownea grandiceps* Jacq. | 27.75 | 13.25 | 273283 | Valley |
| YNP126 | *Brownea grandiceps* Jacq. | 34 | 7 | 342139 | Valley |
| YNP127 | *Brownea grandiceps* Jacq. | 16 | 23 | 165796 | Ridge |
| YNP128 | *Brownea grandiceps* Jacq. | 1 | 12 | - | Valley |
| YNP130 | *Brownea grandiceps* Jacq. | 8.75 | 14.25 | 86695 | Ridge |
| YNP131 | *Brownea jaramilloi* Á.J. Pérez & Klitg. | 27 | 17 | - | Ridge |
| YNP132 | *Brownea jaramilloi* Á.J. Pérez & Klitg. | 27.75 | 17.5 | 270478 | Ridge |
| YNP133 | *Brownea jaramilloi* Á.J. Pérez & Klitg. | 28.25 | 15.25 | 284110 | Ridge |
| YNP134 | *Brownea jaramilloi* Á.J. Pérez & Klitg. | 27.5 | 17.5 | 274514 | Ridge |
| YNP135 | *Brownea jaramilloi* Á.J. Pérez & Klitg. | 28.75 | 15.75 | - | Ridge |
| YNP136 | *Brownea grandiceps* Jacq. | 28 | 15 | 284084 | Ridge |
| YNP137 | *Brownea jaramilloi* Á.J. Pérez & Klitg. | 28.25 | 15 | 284136 | Ridge |
| YNP138 | *Brownea grandiceps* Jacq. | 34.25 | 7.5 | 342301 | Valley |
| YNP139 | *Brownea grandiceps* Jacq. | 34.5 | 7.75 | 342275 | Valley |
| YNP140 | *Brownea jaramilloi* Á.J. Pérez & Klitg. | 27 | 17.5 | 270498 | Ridge |
| YNP141 | *Brownea grandiceps* Jacq. | 34.25 | 7.5 | 342298 | Valley |
| YNP142 | *Brownea jaramilloi* Á.J. Pérez & Klitg. | 17.25 | 18.25 | 174840 | Ridge |
| YNP143 | *Brownea grandiceps* Jacq. | 34.25 | 7.75 | 342311 | Valley |
| YNP144 | *Brownea grandiceps* Jacq. | 8.75 | 14.5 | 83325 | Ridge |
| YNP145 | *Brownea jaramilloi* Á.J. Pérez & Klitg. | 44.5 | 23.25 | - | Ridge |
| YNP146 | *Brownea grandiceps* Jacq. | 9.25 | 12.25 | - | Valley |
| YNP147 | *Brownea grandiceps* Jacq. | 9.75 | 12.5 | 92959 | Valley |
| YNP148 | *Brownea grandiceps* Jacq. | 34 | 7 | 342114 | Valley |
| YNP149 | *Brownea grandiceps* Jacq. | 39.25 | 16.5 | - | swamp |
| YNP150 | *Brownea grandiceps* Jacq. | 44.25 | 10.75 | 440330 | Ridge |
| YNP151 | *Brownea grandiceps* Jacq. | 17 | 21.75 | 175820 | Ridge |
| YNP153 | *Brownea grandiceps* Jacq. | 39 | 16.25 | 390667 | swamp |
| YNP154 | *Brownea grandiceps* Jacq. | 33.75 | 23.5 | - | Ridge |
| YNP155 | *Brownea grandiceps* Jacq. | 49 | 15.75 | 494296 | Valley |
| YNP156 | *Brownea grandiceps* Jacq. | 49.5 | 15.5 | 494174 | Valley |
| YNP157 | *Brownea grandiceps* Jacq. | 39.75 | 16.5 | 394144 | swamp |

| | | | | | |
|---|---|---|---|---|---|
| YNP158 | *Brownea grandiceps* Jacq. | 46.75 | 22 | 465602 | Ridge |
| YNP159 | *Brownea jaramilloi* Á.J. Pérez & Klitg. | 12.25 | 15.75 | 123676 | Ridge |
| YNP160 | *Brownea grandiceps* Jacq. | 39.75 | 16.5 | 394121 | swamp |
| YNP161 | *Brownea grandiceps* Jacq. | 39 | 16 | 394052 | swamp |
| YNP162 | *Brownea grandiceps* Jacq. | 48 | 15.25 | 484468 | Valley |
| YNP163 | *Brownea grandiceps* Jacq. | 17.5 | 21.25 | 175703 | Ridge |
| YNP164 | *Brownea grandiceps* Jacq. | 39.25 | 16.75 | 394164 | swamp |
| YNP165 | *Brownea grandiceps* Jacq. | 39.75 | 16 | 394116 | swamp |
| YNP166 | *Brownea grandiceps* Jacq. | 10.25 | 24.75 | 105976 | Ridge |
| YNP167 | *Brownea grandiceps* Jacq. | 33.25 | 23.5 | 335326 | Ridge |
| YNP168 | *Brownea grandiceps* Jacq. | 33.25 | 23.25 | 335207 | Ridge |
| YNP169 | *Brownea grandiceps* Jacq. | 17.5 | 21.25 | 175698 | Ridge |
| YNP170 | *Brownea grandiceps* Jacq. | 17 | 21.25 | 175630 | Ridge |
| YNP171 | *Brownea grandiceps* Jacq. | 49.75 | 24 | 496303 | Ridge |
| YNP172* | *Brownea* "rosada" Jacq. | 48.75 | 25.75 | 480587 | Ridge |
| YNP173* | *Brownea* "rosada" Jacq. | 48 | 23 | 480580 | Ridge |
| YNP174* | *Brownea jaramilloi* Á.J. Pérez & Klitg. | 5 | 19 | 55158 | Ridge |
| YNP175* | *Brownea jaramilloi* Á.J. Pérez & Klitg. | 5 | 20 | 55449 | Ridge |
| YNP176* | *Brownea jaramilloi* Á.J. Pérez & Klitg. | 7 | 8 | 72048 | Ridge |
| YNP177* | *Brownea jaramilloi* Á.J. Pérez & Klitg. | 2.5 | 2.25 | 6032 | Ridge |
| YNP178* | *Brownea jaramilloi* Á.J. Pérez & Klitg. | 48 | 23 | 480588 | Ridge |
| YNP180* | *Brownea grandiceps* Jacq. | 50 | 4 | 50964 | Ridge |

MOLECULAR ECOLOGY

**Table S3** Population genetic statistics for the dataset containing all SNPs for all loci, calculated using the Stacks *populations* module. Statistics are shown for each 'population', comprised of *B. grandiceps* (including the individuals identified as *B.* "rosada") and *B. jaramilloi*. The first five rows in the table display descriptive statistics, namely the number of individuals in the dataset, the percentage of polymorphic loci, the number of non-shared ('private') alleles, the total number of nucleotide sites in the and the total number of variant nucleotide sites (including the variance and standard error calculated from this value). The subsequent rows show population genetic statistics, along with their variance and standard error. These are: total heterozygosity, homozygosity, π (nucleotide diversity, i.e. the degree of polymorphism in the population) and $F_{is}$ (inbreeding coefficient, i.e. the proportion of polymorphisms in the population present in a single individual).

| | ***Brownea grandiceps* (incl. *B.* "rosada")** | ***Brownea jaramilloi*** |
|---|---|---|
| **Number of individuals** | 131 | 40 |
| **% Polymorphic loci** | 1.83435 | 1.78222 |
| **Number of private alleles** | 7,579 | 747 |
| **Total number of sites** | 4,320,012 | 5,351,291 |
| **Total number of variant sites** | 79,991 | 102,951 |
| *Variance* | 525.71544 | 48.45287 |
| *Standard Error* | 0.01102 | 0.00301 |
| **Heterozygosity** | 0.00279 | 0.00261 |
| *Variance* | 0.00075 | 0.00075 |
| *Standard Error* | 0.00001 | 0.00001 |
| **Homozygosity** | 0.99721 | 0.99739 |
| *Variance* | 0.00075 | 0.00075 |
| *Standard Error* | 0.00001 | 0.00001 |
| **π** | 0.00453 | 0.00475 |
| *Variance* | 0.0015 | 0.00166 |
| *Standard Error* | 0.00002 | 0.00002 |
| **$F_{is}$** | 0.00716 | 0.00766 |
| *Variance* | 0.00487 | 0.00607 |
| *Standard Error* | 0.01103 | 0.00301 |

MOLECULAR ECOLOGY

**Methods S1** Supplementary methods for the phylogenomics section of the study across *Brownea*.

### *Phylogenomic Taxon Sampling*

A species list was compiled containing all species within the Brownea clade using the Plant List (http://www.theplantlist.org/), Tropicos (http://www.tropicos.org/) and generic monographs (Cowan 1953; Klitgaard 1991; Redden, Herendeen, & Lewis 2018). This was to ensure that taxonomically accepted species were included in analyses. Additionally, the voucher specimens were examined, and their determination updated as appropriate. Sampling was targeted to include specimens from throughout the geographical ranges of as many species as possible, which were represented by multiple accessions from throughout their range where possible. Accessions were acquired from both silica material and herbarium specimens collected from the following herbaria: AAU, E, K, NY, US. This investigation utilizes the 'lineage' species concept (as defined by De Quieroz (2007)), since rainforest tree species act as independently-evolving metapopulations through effective gene flow despite the fact that non-monophyly is common in these taxa for many gene regions due to a large $N_e$ and long generation times (Pennington & Lavin 2016).

### *Phylogenomic Library Preparation and Sequencing*

DNA extractions were carried out using 20 mg of dried leaf material with the CTAB method (Doyle & Doyle 1987). DNA concentrations were measured using a Quantus fluorometer (Promega, Wisconsin, USA).

Libraries were prepared using the NEBNext® Ultra™ II DNA Kit (New England Biolabs, Massachusetts, USA) with a modified protocol to account for fragmented DNA. Samples with high molecular weight DNA (>1000 bp, measured with a TapeStation 4200 (Agilent Technologies, California, USA)) were sheared to around 600bp with a Covaris focussed ultrasonicator M220 (Covaris, Massachusetts, USA). Following this, end-preparation and Illumina adaptor ligation were undertaken according to the NEBNext protocol. In order to prevent loss of DNA during library preparation, samples with low molecular weight were excluded from pre-PCR size selection step suggested by the NEBNext kit. However, a post-ligation clean-up was performed using 0.9x Agencourt Ampure XP magnetic beads (Beckman Coulter, California, USA). Libraries containing a range of insert sizes were then amplified using PCR, following the protocol outlined in Table SM1.1.

| Cycle step | Temperature | Time | Nº Cycles |
|---|---|---|---|
| *Initial Denaturation* | 98°C | 30 seconds | 1 |
| *Denaturation* | 98°C | 10 seconds | 12 |
| *Annealing/Extension* | 65°C | 75 seconds | |
| *Final Extension* | 65°C | 5 minutes | 1 |
| *Hold* | 4°C | ∞ | - |

**Table SM1.1:** Amplification conditions used in the NEBNext ® Ultra™ II protocol to amplify adaptor-ligated DNA libraries.

The amplified libraries were subsequently size-selected using 0.7x Agencourt Ampure XP magnetic beads in a two-step process (i.e., a size-selection using 0.4x Ampure XP followed by one using 0.3x Ampure XP), aiming for a ~600 bp fragment length, which was verified using a TapeStation. Following this, all samples were normalized to 20nM. Because unique adapter set combinations were used all 73 samples were pooled at equimolar levels, resulting in a 20nM library pool.

Hybrid bait capture was performed using the MyBaits v3.02 protocol (Arbor Biosciences, Michigan, USA) for samples S1-S48 and the MyBaits v4.01 protocol for S49-S105. This method enriches target gene regions to allow the sequencing of degraded DNA samples (such as those from museums or herbaria). The bait set used for hybrid capture (manufactured by MYcroarray (Michigan, U.S.A.)) targeted 289 nuclear genes and was designed for the legume subfamily Detarioideae (Ojeda *et al.*, 2019). This bait set was designed using transcriptome sequences of four species representing the major clades within the Detarioideae (de la Estrella *et al.*, 2018). These four transcriptomes were used to identify open reading frames from which coding regions with at least 100 amino acids were extracted. In order to avoid including paralogous regions in this bait set, a CD-HIT (W. Li & Godzik 2006) search was performed to find coding regions present as single or low-copy genes in the genomes of six other legume species. A self-BLAST search (Gish & States 1993) was performed on these genes to discard regions with multiple hits, which were taken to be potential paralogs. Orthologs were identified among the coding regions using a BlastP (Gish & States 1993), and only those orthologs which showed a topology congruent previous work (de la Estrella *et al.*, 2017; de la Estrella *et al.*, 2018) were further refined and manufactured as RNA baits. Hybridization was performed for 30 hours at 65 °C, followed by three washes using Dynabeads® MyOne™ Streptavidin C1 Beads (Thermo Scientific, Massachusetts, USA). Enriched libraries were eluted, cleaned with 0.7x Agencourt Ampure XP beads and amplified using 14 cycles of PCR. The DNA fragment length distribution was measured using a TapeStation and DNA concentration of enriched libraries was measured using a Quantus. The final library pools containing samples S1-S48 were sequenced on the Illumina MiSeq platform (Illumina, San-Diego, USA) with a 2x300bp paired-end run at RBG Kew, and S49-S105 were sequenced with a 2x150bp paired-end run on the HiSeq platform by Macrogen Inc. (Seoul, South Korea).

### *Quality-filtering, Read Assembly and Alignment*

Illumina sequencing reads were quality-checked using their phred-33 score in the program *FASTQC* v0.11.3 (Andrews 2010). Following quality-checking, reads had their adapter sequences trimmed using *Trimmomatic* v.0.3.6 (Bolger, Lohse, & Usadel 2014), permitting a maximum of four mismatches, a palindrome clip threshold of 30 and a simple clip threshold of 6. Reads were subsequently quality-filtered in *Trimmomatic* by removing bases with a phred-33 score <28 from the beginning and end of the reads, and by using a four-base sliding window to remove bases with an average phredd-33 score of <30. In addition, reads shorter than 36 bases long were removed from the dataset.

After quality-filtering, loci were assembled with the HybPiper pipeline v1.2 (Johnson *et al.*, 2016). Using this pipeline, reads were mapped to the target genes used to design the Detarioideae bait kit with the BWA algorithm (H. Li & Durbin 2009), after which they were assembled into contigs representing coding gene regions with *SPAdes* v3.11.1 (Bankevich *et al.*, 2012) using the default settings for the program, except for an increased minimum coverage cut-off of 8x rather than 5x. The coding sequences for each individual were extracted by assembling contigs to their corresponding target gene regions with *Exonerate* (Slater & Birney 2005), which is part of the HybPiper pipeline. This program chooses the best contig for each gene region based on its percentage identity (threshold = 55%), coverage depth (threshold = 10x) and length relative to the target gene sequence (threshold = 90%). The gene recovery for each region was then assessed by examining read lengths and the number of reads mapped per individual.

Potentially paralogous loci were identified using the Python (Python Software Foundation 2010) script ‘*paralog_investigator.py*’, which is distributed with the HybPiper pipeline. This script identifies alternative (i.e., potentially paralogous) contigs associated with each gene region for each individual, allowing these regions to be excluded from the dataset before alignment. Following this, the coding sequences extracted by *Exonerate* were aligned by gene region (excluding those with potential paralogs) using 1,000 iterations in *MAFFT* v7.215 (Katoh & Standley 2013) and the ‘*—adjustdirectionaccurately’* option to incorporate reversed sequences. These alignments were then visually inspected for poor sequence quality using *Geneious* v. 8.1.9 (https://www.Geneious.com) in order to remove taxa with mostly missing data and to prevent the inclusion of poorly-recovered loci into the dataset.

### *Inferring phylogenetic networks*

In order to infer phylogenetic networks from the 220 single-accession-per-lineage gene trees, the function ‘*readTrees2CF*’ was used to calculate concordance factors (CF) in order to estimate the proportion of gene trees which support each possible relationship between quartets of taxa. *SNaQ!* was subsequently used to calculate the negative log (-log) pseudolikelihood of a tree under ILS with no hybridization ($h$ = 0), as well as the -log pseudolikelihoods of networks representing an increasing number of hybridization events ($h$ = 1 to 4). Pseudolikelihoods in a phylogenetic network are approximated from the likelihood formulas of its four-taxon subnetworks, and as such the calculated likelihoods are not independent (hence ‘pseudo’-likelihood). However, this also means that they are less computationally complex to calculate (Liu, Yu, & Edwards 2010; Solís-Lemus, Bastide, & Ané 2017). *SNaQ!* analysis was performed using 10 independent runs for each value of $h$ to ensure convergence, with the single-accession *ASTRAL* species tree as a starting tree. Different values of $h$ were then compared to ascertain the number of hybridization events which caused the greatest increase in -log pseudolikelihood. This is because -log pseudolikelihoods are expected to increase rapidly with an increasing number of hybridization events until the optimum is reached, after which the increase occurs more slowly as the number of hybridization events increases (Solís-Lemus & Ané 2016). Pseudolikelihood scores for each value of $h$ were plotted as a line graph in R v.3.4.4 (R Development Core Team 2013), and the network with the highest -log pseudolikelihood was visualised using *PhyloNetworks* v0.11.0.

Following network estimation, the fit of the CFs generated from the gene trees were compared both to those expected under a model including only ILS (i.e., a ‘tree-like’ model, $h$ = 0) and a model incorporating hybridization (i.e., a phylogenetic network, $h$ > 0). This was done using the Tree Incongruence Checking in R (TICR) test (Stenz *et al.*, 2015) implemented in the R package *PHYLOLM* (Ho & Ané 2014). The TICR test measures the goodness-of-fit of the observed CFs to the coalescent model to ascertain whether ILS explains most of the gene tree discordance observed. If the coalescent model does not explain the gene tree discordance adequately, then the network-like model is more likely to explain the discordance. In order to test this hypothesis, the distributions of the $P$-values describing the goodness-of-fit for each quartet (which consists of $P$-values binned into four categories: ($P$ < 0.01, 0.01–0.05, 0.05–0.1, 0.1–1)) are compared with the distribution of $P$-values expected under the coalescent model. A $X^2$ test is performed on these distributions, and if $P \leq 0.05$ for the $X^2$ test, then the observed CF values do not fit the coalescent model. In other words, if there are more $P$-values <0.05 than would be expected by chance, then the data are not tree-like. This was performed using the function ‘*test.one.species.tree*’ in *PHYLOLM.*

MOLECULAR ECOLOGY

**Methods S2** Methods for the population genomic section of the study, focussed on two co-occurring *Brownea* species in Yasuní National Park, Ecuador.

### Study site

Yasuní National Park (YNP) is an area of lowland tropical rainforest covering around 1.6 million hectares of the Ecuadorian Amazon in north-western South America. The majority of the park comprises unbroken primary rainforest, with a very high alpha-diversity of tree species (Bass *et al.*, 2010; Valencia, Balslev, & Miño 1994). Within YNP is a 50ha permanent forest plot, the south-western edge of which is located at 0° 41' 0.5" S latitude, 76° 23' 58.9" W longitude, and the altitude above sea level ranges between 215-248m. The 50ha plot is subdivided into 1,250 20m x 20m subplots, and every tree above 1cm diameter-at-breast-height has been identified and labelled with a unique identifying tag. All trees sampled for this study were taken from within this 50ha plot. The location of Yasuní National Park, as well as an approximate range map for *B. grandiceps* and *B. jaramilloi* and the locations of georeferenced collections are shown in Figure SM2.1. In order to draw this map georeferenced collections were downloaded from GBIF (www.GBIF.org, 3rd October 2018: GBIF Occurrence Downloads https://doi.org/10.15468/dl.vgy5oz & https://doi.org/10.15468/dl.yfqlkj) and cross-referenced using the BIEN portal (http://www.biendata.org), as well as taxonomic treatises on the two species (Klitgaard 1991; Pérez *et al.*, 2013). Maps were then plotted using QGIS v. 3.2.0 (QGIS Development Team 2017).

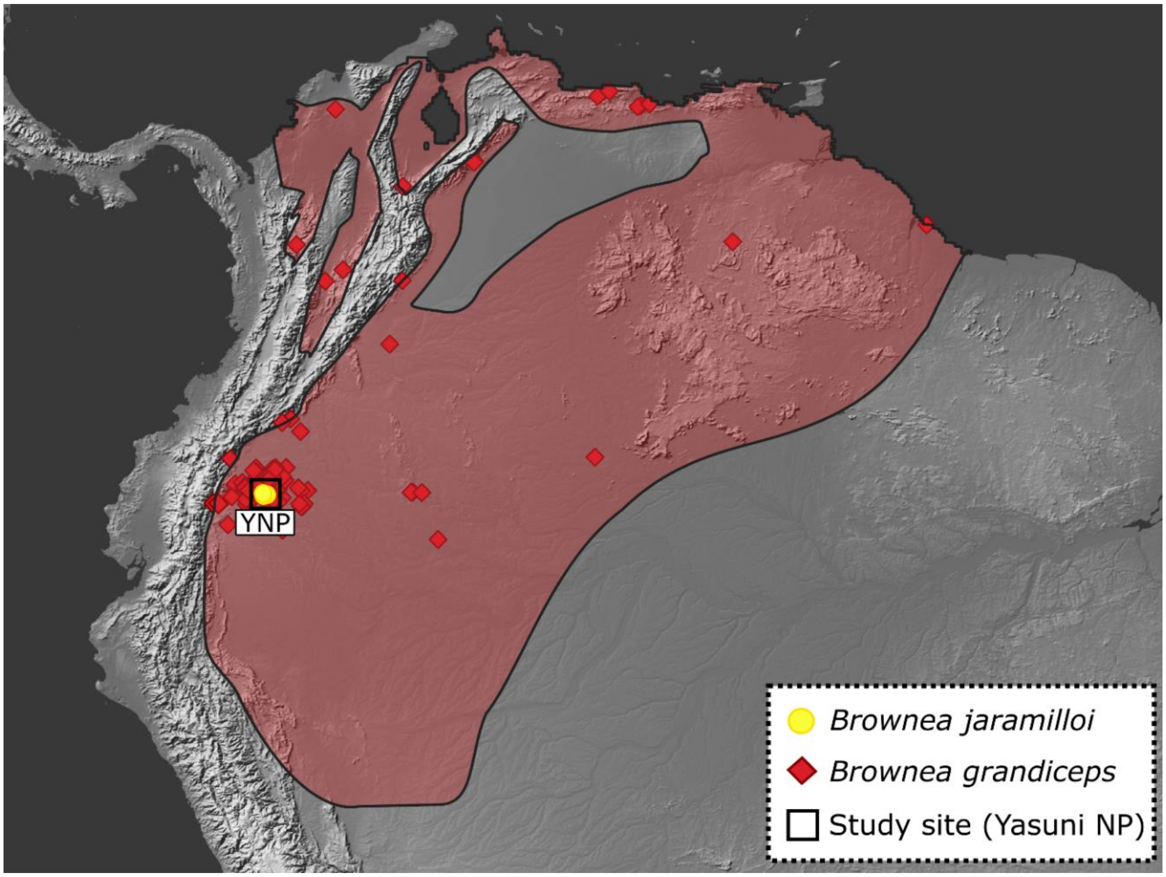

*Figure SM2.1*: Map of South America, showing the sampling location for all accessions examined during the population genomic study (Yasuní National Park) and approximate distributions of *Brownea grandiceps* (in red) and *B. jaramilloi* (in yellow), which appears to be endemic to the Ecuadorian Amazon. The red and yellow points are refined georeferenced collections collected from GBIF (www.gbif.org) and were plotted on the map of South America in QGIS v.3.2.0.

### *Sampling*

One-hundred and seventy-one individuals belonging to two species (*B. grandiceps* and *B. jaramilloi*), as well as a putative hybrid between the two species (*B.* "rosada"), were sampled for this study. In total, 128 individuals of *B. grandiceps*, 40 individuals of *B. jaramilloi* and three individuals of *B.* "rosada" were genotyped, representing their relative proportions within the Yasuní 50ha plot. Of these accessions, 162 were collected from the 50ha forest plot in October-December 2016, and the remaining nine were silica collections made in 2002 from the same plot.  Specimens were identified to species level based on leaf characters and inflorescences found on the specimens collected from the field, which were then cross-referenced with the Yasuní plot census list.

In order to sample individuals from the 50ha plot, 32 of the total 1,250 20x20m sub plots were randomly selected. Sampling was stratified by defining two separate subplot types- 'valley' and 'ridge', in order to sample a representative number of individuals from both species. This is because *B. jaramilloi* tends to favour the 'ridge' subplot type. Using the existing elevation cut-offs by which the plot is stratified, 'valley' subplots were defined as having a median elevation ≤227.2 m, whereas 'ridge' subplots were defined as having a median elevation >227.2 m (Valencia *et al.*, 2004). As such, 16 subplots of each subplot type were randomly selected, from which accessions were collected (shown in Figure SM2.2).

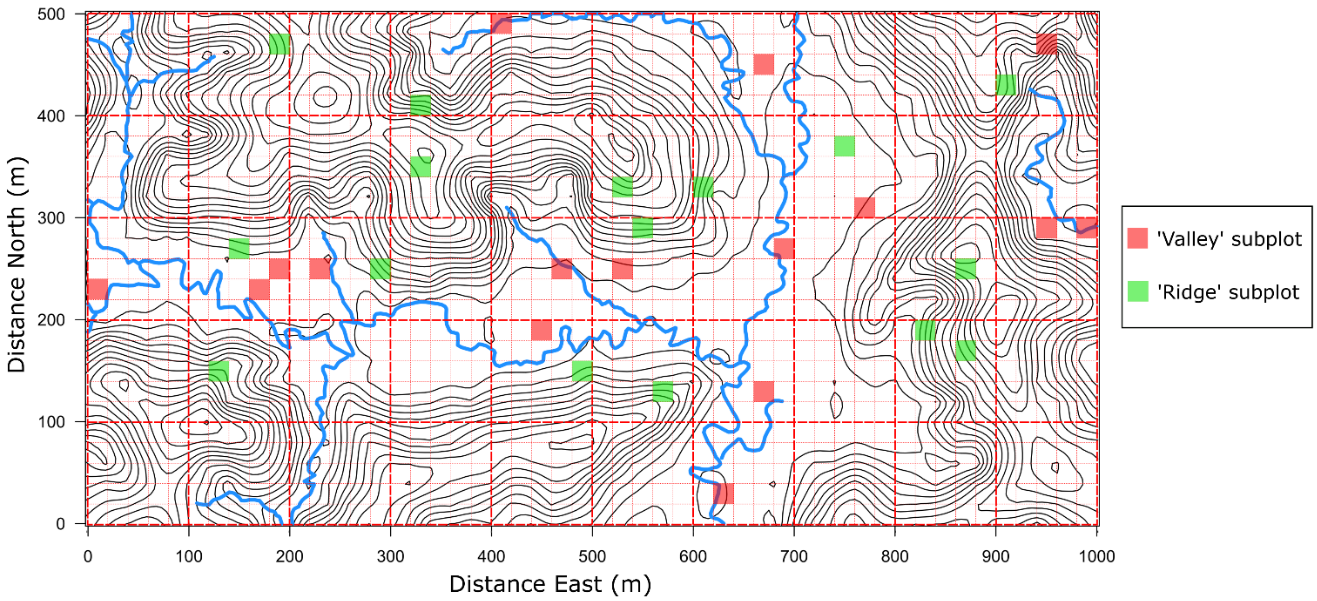


*Figure SM2.2:* 20x20m subplots sampled in Yasuní National Park, Ecuador. There are 16 subplots belonging to two different types: 'Valley' (mean elevation ≤227.2 m) and 'Ridge' (mean elevation >227.2 m) (Valencia *et al.*, 2004). Dashed gridlines denote 20x20m plots within the entire 50ha forest plot.

Leaf material was collected from every individual belonging to both *Brownea* species and their hybrid encountered within each subplot, both from mature trees (marked with tags from the recent plot census) as well as from juvenile trees (i.e. those under 1 cm diameter-at-breast-height (DBH) and lacking a census tag). Collected leaf material was pressed and dried using an herbarium press and drying cabinet. Specimens were dried for a maximum of 12 hours in order to minimize damage to genomic DNA.

### *Library preparation and sequencing*

Genomic DNA was extracted from 30 mg of lyophilized leaf tissue using a modified CTAB protocol (Doyle & Doyle 1987), after which extracts were cleaned using a QIAGEN plant minikit (QIAGEN, Hilden, Germany) column cleaning stage. RNA was removed from extracts using RNase A, as per the manufacturers protocol. Following this, the DNA concentration of the samples was measured using a Quantus fluorometer (Promega, Wisconsin, USA).

Libraries were prepared using 500ng of DNA template from each sample, which was digested for three hours using the restriction enzymes *EcoRI* and *mspI* (New England BioLabs, Massachusetts, USA), in accordance with the ddRADseq protocol (Peterson *et al.,* 2012). Following digestion, samples were ligated to universal Illumina P2 adapters, and barcoded using 48 unique Illumina P1 adapters. Samples were pooled in sets of 48 at equimolar concentrations following normalisation. In total, four ddRAD libraries of 48 samples each were prepared, which were then size-selected to between 375-550 bp using a Pippinprep electrophoresis machine (Sage Science, Massachusetts, USA). Libraries were amplified using 12 cycles of PCR with Phusion High-Fidelity DNA polymerase (New England Biolabs, Massachusetts, USA), after which they were multiplexed, with a specific multiplexing index being used for each library pool of 48 samples. Reactions were cleaned between each step in the ddRAD protocol using Agencourt Ampure XP magnetic beads (Beckman Coulter, California, USA), and quantified using a Quantus fluorometer. Sequencing was performed using a single-lane, paired-end 150bp run on the HiSeq 3/4000 platform, undertaken at the Norwegian Sequencing Centre, Oslo (www.sequencing.uio.no).

### *De-novo read assembly*

Paired-end Illumina reads were processed into loci *de novo* using the Stacks pipeline v2.1 (Catchen *et al.,* 2011). First, raw Illumina reads were de-multiplexed and quality-filtered by removing reads with an average Phred score <10 using the '*process_radtags*' function in Stacks. Following this, the *denovo_map.pl* program in Stacks was run, which first creates 'stacks' of reads for each individual, from which RAD tags (i.e. 'loci') are inferred and SNPs identified (*ustacks*). These stacks are subsequently combined for each locus across all individuals (with each allele merged) to create a consensus catalogue of alleles from the sampled populations (*cstacks*). Finally, every locus from each individual is matched to the catalogue generated by *cstacks* in order to identify which SNP combinations are present in the populations under study (*sstacks*). A minimum coverage depth (-*m*) of three was used to create each stack, with a maximum within-individual mismatch distance (-*M*) of seven nucleotides- this value was particularly high to account for possible polyploidy of hybrids. In order to process an acceptable number of loci from individuals belonging to both species and their hybrids, a mismatch distance (-*n*) of seven nucleotides was used by *cstacks* to stack loci. Model parameters for the various programs within the Stacks pipeline were chosen using the recommendations in Paris *et al*. (2017). In addition, numbers of loci retained under different combinations of the *m* and *n* parameters ($m$=2, $n$= 3; $m$=3, $n$=2; $m$=3, $n$=4; $m$=4, $n$=5) were compared based on a random subset of the ddRADseq data, which made minimal changes to the results. After the initial single-end loci were constructed by *ustacks*, *cstacks* and *sstacks* the *tsv2bam* module was run, which incorporates the reverse Illumina reads, allowing as much of the Illumina output to be used as is possible. Individuals with sequencing coverage under 7.5x were removed after processing. Data were further filtered using the *'populations*' module of Stacks: only loci found in ≥40% of individuals of both *Brownea* species and their hybrids were retained, and minor alleles that were present in <5% of individuals were excluded from the dataset to control for genotyping error. This resulted in a dataset containing 22,046 loci with 120,085 SNPs for 171 individuals. Population genetic statistics were calculated by the *populations* module for each of the two species (including the number of variant sites, heterozygosity, homozygosity, $\pi$ and $F_{is}$). In addition to this, the *populations* module was used to extract a dataset containing only one SNP per locus with the '*—write_single_SNP'* option. This reduces the likelihood of using linked alleles, which is necessary to prevent linkage disequilibrium (LD) affecting parameter estimates in programs that assume no LD.  The first SNP of each locus was used because it is likely to have lower sequencing error associated with it, given that sequencing errors rates tend to increase towards the end of a read. This process resulted in a dataset containing 19,130 loci with 19,130 SNPs for 171 individuals.

### *Gene flow and population structure*

The full dataset containing all SNPs per RAD locus was visualised using a principal component analysis performed using the R function *prcomp*, and plotted with the package *ggplot2* (Wickham 2016) in R version 3.5.1 (R Development Core Team 2013). In addition to this, a Neighbour net plot was inferred with uncorrelated P-distances in the program *SPLITSTREE* v4.14.6(Huson & Bryant 2005).

Population structure was estimated and visualised in the program *fastSTRUCTURE* v1.0 (Raj, Stephens, & Pritchard 2014) using the single-SNP-per-locus dataset in order to account for linkage between SNP loci, as per the assumptions of the program. The number of populations (*K*) was selected based on the value with the largest difference in marginal likelihood, ascertained after testing multiple *K* values (between 1-5). *fastSTRUCTURE* was run again using 40 individuals of each of the two species (*B. grandiceps* and *B. jaramilloi*) to test whether the same value of *K* was inferred, and that the inferred patterns of introgression were not biased by different population sizes.

The command-line version of the program *NEWHYBRIDS* v1.1 (Anderson and Thompson (2002), https://github.com/eriqande/newhybrids) was used to categorize genotyped individuals into different hybrid classes (pure, F1, F2, and backcrosses). Five hundred loci were randomly subsampled from the single-SNP-per-locus dataset and were used for the analysis, due to the computational limitations of the program. Loci were subsampled for 169 of 171 individuals, the missing two of which were removed due to the amount of missing data they possessed. *NEWHYBRIDS* was run using 50,000 MCMC sweeps with 50,000 burn-in sweeps, and the Jeffries-like priors were used for both the allele frequency ($\theta$) and mixing proportion ($\pi$) parameters because no prior information was available regarding allele frequencies or the amount of admixture. No hybrid classification was given to individuals *a priori* in order to reduce bias, as well as to determine whether the SNPs used represented the whole dataset through their correct identification of pure individuals. Three analyses were run on three different 500-SNP subsets, which delivered convergent results.

In order to quantify the dynamics of introgression at each locus within the *Brownea* lineages and their hybrids, Bayesian estimation of genomic clines (*bgc*) v1.03 (Gompert & Buerkle 2011) was used. This program creates a genomic admixture gradient (or 'cline') for all individuals, ranging from one species to the other. From this, the program compares the probability of ancestry at a locus (φ) to an individual's genome-wide average ancestry (*h*). *bgc* estimates two parameters for each locus: 'α' and 'β'. α is the position of the genomic cline's centre relative to *h* = 0.5. Positive values of α indicate an increased probability of ancestry from one parent at a specific locus when compared to the genome-wide average, and negative values indicate an increased probability of ancestry from the other parent at a specific locus. If α is equal to zero, then there is the same probability of ancestry from each parent as the genome-wide average. Therefore, the α parameter can be approximated as a measure of the 'direction' of introgression. The β parameter represents the genomic cline's gradient and is the rate of transition in the probability of ancestry at a locus when compared to the genome-wide average. Positive values of β are indicative of a rapid rate of change in the probability of ancestry relative to the genome-wide average, which occurs with reduced introgression (i.e., selection against hybrid genotypes). Negative values of β indicate a more gradual change in the probability of ancestry due to more shared alleles resulting from greater introgression. If β equals zero, then the gradient of the cline is φ = *h*, which is the null expectation. In other words, if φ = *h* the locus introgresses at the genome-wide average rate, as might be expected when there is no selection occurring. As such, β can be thought of as the 'resistance' or 'receptivity' to introgression for a specific locus (Gompert & Buerkle 2011; Gompert, Parchman, & Buerkle 2012). A diagram of hypothetical genomic clines is shown in Figure SM2.3.

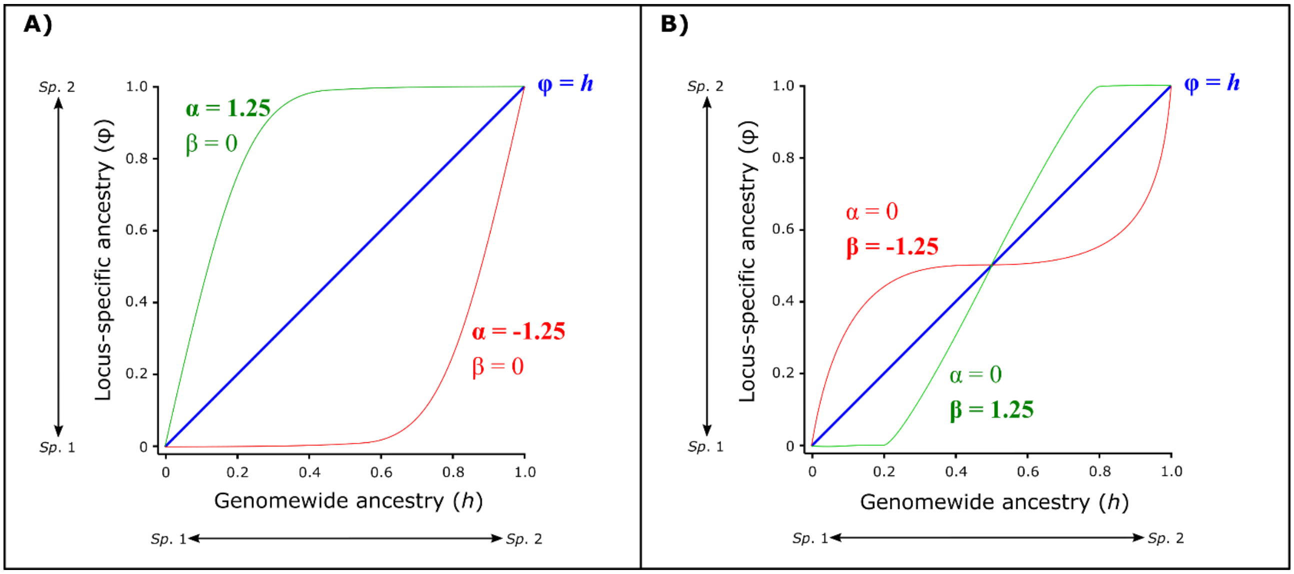


*Figure SM2.3:* Hypothetical genomic clines to illustrate what the parameters α and β correspond to. Each line (excluding φ = *h*, which is the null expectation) represents a locus within the two hybridizing populations, and the shape of the cline represents the relationship between φ and *h* in all individuals within the populations at that locus. Genome-wide ancestry (*h*) is the proportion of an individual's genome inherited from one parental species, and φ is the probability of locus-specific ancestry from the same parental species (Gompert & Buerkle 2012; Gompert & Buerkle 2011; Lexer *et al.*, 2007). The left pane (**A**) shows a hypothetical locus with more introgression from species 1 (green line) and another locus with more introgression from species 2 (red line). The right pane (**B**) shows a hypothetical locus that is very resistant to introgression (green line) and another locus that is very prone to introgression (red line).

In order to estimate these parameters from the single-SNP dataset containing 19,130 loci, two runs of 50,000 MCMC steps were undertaken in the program *bgc* (Gompert & Buerkle 2012), including 25,000 steps discarded as burn-in. Genomic data were converted from the *genepop* format (Raymond 1995;

Rousset 2008) to *bgc* input format using the R package *genepopedit* (Stanley *et al.*, 2017) in R v. 3.5.1. The posterior distribution was thinned every 10 steps, resulting in 5000 samples per locus for parameter inference. Parental and admixed populations were defined according to the mean *Q* values from *fastSTRUCTURE*, where *Q* represents the genetic admixture proportion of an individual. Since there were two parental populations, individuals with mean *Q* >0.08 and <0.92 were defined as 'admixed'. A mean *Q* of ~0 corresponds to *B. grandiceps*, and a mean *Q* of ~1 denotes *B. jaramilloi*. The MCMC runs were checked for convergence using the log-likelihood output of *bgc* in Tracer v1.6 (Rambaut *et al.*, 2015) and with the R package *coda* (Plummer *et al.*, 2006) using Geweke's diagnostic (Geweke 1991). Geweke's diagnostic outputs a *Z*-value to test for equality of means between the start and end of MCMC runs, excluding burn-in. Estimates of the α and β parameters and their 99% posterior probability credible intervals were generated from the *bgc* output using the program *estpost* (Jann 2017), after which loci with 'excess ancestry' were identified by filtering out all loci whose 99% credible intervals included zero, thereby attaining all positive and negative non-zero estimates of the α and β parameters. Finally, statistically extreme 'introgression outliers' were identified for both parameters by identifying loci whose median estimates were not included in the 99% posterior probability credible intervals (Gompert & Buerkle 2011).
